# Massively Parallel FPGA Hardware for Spike-By-Spike Networks

**DOI:** 10.1101/500280

**Authors:** David Rotermund, Klaus R. Pawelzik

## Abstract

While inspired by the brain, currently successful artificial neural networks lack key features of the biological original. In particular, the deep convolutional networks (DCNs) neither use pulses as signals exchanged among neurons, nor do they include recurrent connections which are both core properties of real neuronal networks. This not only puts to question the relevance of DCNs for explaining information processing in nervous systems but also limits their potential for modeling natural intelligence.

Spike-By-Spike (SbS) networks are a promising new approach that combines the computational power of artificial networks with biological realism. Instead of separate neurons they consist of neuronal populations performing inference. Even though the underlying equations are rather simple implementations of such networks on currently available hardware are several orders of magnitude slower than for comparable non-spiking deep networks.

Here, we develop and investigate a framework for SbS networks on chip. Thanks to the communication via spikes, already moderately sized deep networks based on the SbS approach allows a parallelization into thousands of simple and fully independent computational cores. We demonstrate the feasibility of our design on a Xilinx Virtex 6 FPGA while avoiding proprietary cores (except block memory) that cannot be realized on a custom-designed ASIC. We present memory access optimized circuits for updating the internal variables of the neurons based on incoming spikes as well as for learning the connection’s strength. The optimized computational circuits as well as the representation of variables fully exploit the non-negative properties of all data in the SbS approach. We compare the sizes of the arising circuits for floating and fixed point numbers. In addition we show how to minimize the number of components that are required for the computational cores by reusing their components for different functions.

## 1 INTRODUCTION

Nowadays, deep neuronal networks (Schmidhuber, 2015) are a basis for successfully applying neuronal networks on problems from artificial intelligence research (Azkarate Saiz, 2015; Silver et al., 2016; Guo et al., 2016; Gatys et al., 2016). The revival of using neuronal networks was provoked by the increase of computational powers in modern computers and boosted even more through modern 3D graphic cards as well as specialized application specific integrated circuits (ASICs) (Sze et al., 2017; Jouppi et al., 2018) and field programmable gate arrays (FPGAs) (Lacey et al., 2016). The most successful type of networks is based on multilayer perceptrons (Rumelhart et al., 1986; Rosenblatt, 1958) and consists of several so called hidden layers. Typically information is processed and feed forward from one hidden layer to the next, beginning at the input layer and ending at the network’s output layer. In theory such a network is able to calculate arbitrary functions. However, for doing so the weights – describing the connection of elements of one layer to the next – must be learned based on the intended task. During learning these weights, the error between the desired outcome and the actual outcome of the network is minimized.

In the realm of brain research, more detailed and biologically realistic spiking neuron models are used (Maass and Bishop, 2001; Davies et al., 2018; Thakur et al., 2018; Pfeiffer and Pfeil, 2018; Izhikevich, 2004) which require a vast amount of computational power (Izhikevich, 2004). There is a large engineering community that constructs neuromorphic hardware for accelerating the necessary computation for simulating these type of neurons, e.g. (Thakur et al., 2018; Davies et al., 2018; Furber et al., 2014; Moore et al., 2012; Wang and van Schaik, 2018). Also, networks in the brain have no simple feed-forward architecture but instead recurrent connections are ubiquitous implying that real information processing is dynamic.

In (Ernst et al., 2007) we presented a different type of neuronal network (called Spike-by-Spike network, SbS) based on the family of generative models (Lee and Seung, 1999, 2001; Salakhutdinov, 2015; Hinton, 2012). Instead of separate neurons they consist of neuronal populations performing inference and the neurons exchange information via stochastic spikes. In terms of computational requirements the SbS network lies in between the traditional perceptron based non-spiking networks and typical spike-based networks.

In terms of biological plausibility, the SbS network is placed in between non-spiking networks (e.g. deep convolutional networks) and networks of spiking neurons with realistic models (e.g. leaky integrate-and-fire neurons, IaF neurons). Comparing it with networks of IaF neurons, the SbS network removes large parts of biological plausibility. However, this comes with a reduction in parameters. There is no need to optimize e.g. time parameters, firing thresholds, or membrane constants because there is no real time left in a SbS network. Instead of many (sometimes thousands) computational steps that are required to get the next spike in a IaF population, a SbS IP requires only one update per neuron to determine the next spike. Such an update requires 3*N* multiplications, 2*N* summations, and the inversion of one value if *N* is the number of neurons in the SbS IP. It is also possible to build recurred networks with IPs (Rotermund and Pawelzik, 2019b). This all together allows SbS IPs to be used for building larger models for understanding information processing in the brain.

Furthermore all entities in SbS networks are described by positive numbers which leads to sparse representations Bruckstein et al. (2008). A recent discovery in the field of machine learning is compressed sensing (Candes et al., 2006), which allows to reconstruct underlying causes from incomplete measurements if the underlying causes are sparse. This lead to applications in many fields from reducing measurement time for Magnetic Resonance Imaging (Lustig et al., 2008) to building swarms of robots for efficiently exploring other planets (Wiedemann et al., 2018). Sparseness and its benefits for information processing is also an important topic in brain research (Olshausen and Field, 2006; Spanne and Jörntell, 2015; Ganguli and Sompolinsky, 2012). In the context of this work, the finding that non-negative representations in a network can be sufficient to induce some degree of sparsity is particularly interesting(Ganguli and Sompolinsky, 2010; Bruckstein et al., 2008). In (Rotermund and Pawelzik, 2019a) we extended the old SbS approach for shallow networks to deep networks consisting of large numbers of inference populations (IPs). In (Rotermund and Pawelzik, 2019b) we show how to use bi-directional information exchange between the layers of deep SbS network for biologically realistic learning.

Figure 1 shows an example SbS network architecture for classifying MNIST benchmark handwritten digits. The shown network consists of several layers. Each layer can perform different computations like convolution, pooling or other learned functions. These layers are all constructed from many IPs and all the IPs realize the same algorithm for updating the latent variables based on arriving spikes as well as learning the weights. The spikes in such a network can flow either only forward, from input to output, (Rotermund and Pawelzik, 2019a) or spikes can flow in both direction to e.g. neighboring layers (Rotermund and Pawelzik, 2019b). In this example, spike are only exchanged between IPs from different layers but not between IPs in the same layer. On a more abstract level such a network can be understood as pools of IPs and input populations, an architecture that defines which of these populations are allowed to interact, and spikes traveling between IPs.

**Figure 1.**
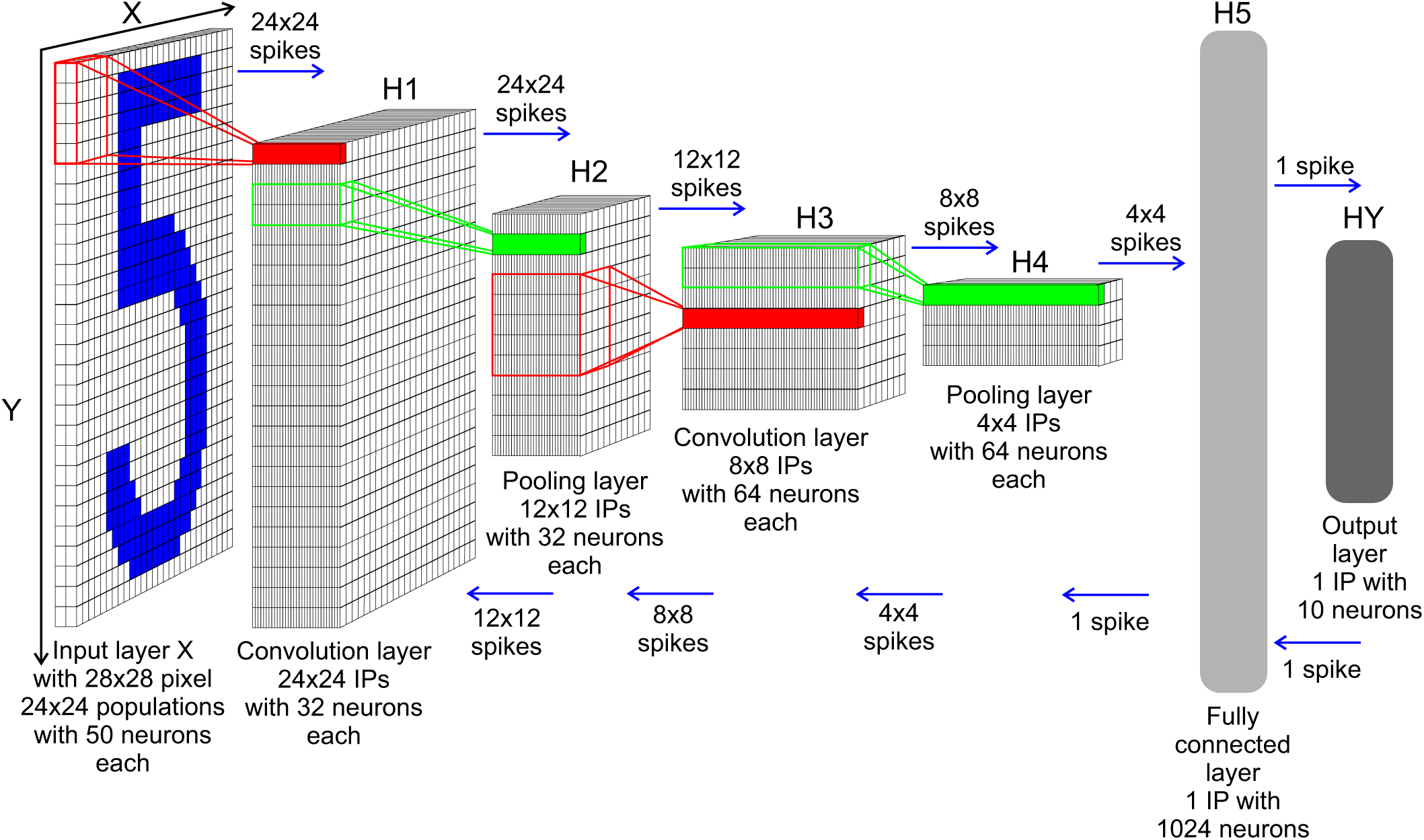
Example SbS network (see (Rotermund and Pawelzik, 2019a) for more details on this network) for analyzing handwritten digits (MNIST benchmark, http://yann.lecun.com/exdb/mnist/). The input image (28×28 gray value pixels) is represented by a 576 node input population and 802 parallel SbS IPs with different number of neurons (with 10, 32, 64, or 1024 neurons per SbS inference population IP, colored columns in layers H1-H4). Thus this network consists of 1378 independent computational elements. The network is organized in one input layer, two convolution layers, two pooling layers, one fully connected layer, and one output layer. All the layers, except the input layer, use the same update dynamic (i.e. updating the internal state of a SbS inference population with an incoming spike). In contrast to usual deep convolutional networks, there is no algorithmic difference for pooling layers, they only have a special arrangement of their weight matrices. Information between elements (SbS inference population and input populations) are exchanged only by spikes. Depending on the architecture of the network, the information can flow either only forward or bi-directionally (as indicated by the blue arrows). The latter can be used to implement local learning rules. In this specific network there are no interactions in between SbS inference populations within the same layer, but in a more complex network they could be helpful.

In the supplemental materials, we have summarized the stylized facts of the SbS algorithm. The important equation for the design of the hardware are recapitulated in section 2.1.

While SbS networks are far less computationally demanding than networks with more detailed spiking neuron models, e.g. leaky integrate-and-fire neurons (IaF), it still requires high computational effort, especially if it is compared to a perceptron-based deep neuronal network. The reduction of the required computing power compared to an IaF is a direct result of time progressing in a SbS network only from one spike to the next. In between spikes, there are no operations happening in a SbS network; formally this time does not exist in such a network. Thus biological realism is traded in for bigger networks that require less computational effort for running them.

However, simulating a SbS network, in particular deep ones with many inference populations for large data sets, is a problem when only normal computer CPUs, GPUs or even computer clusters are available. Optimally simulating a SbS network benefits from a large number of parallel cores with a medium amount of non-shared & directly accessible memory. Every IP in a SbS network is a compact local Module that only communicates via spikes with its environment. This allows to parallelize the whole network into arbitrarily many parallel IPs. In the example of the MNIST network, shown in figure 1, it is best realized by 802 individual threads (In bigger network, this number will significantly increase.). One thread per IP would be the optimal solution. The information processing within IPs can even be asynchronous to the rest of the network as long as the spikes between the populations can be exchanged successfully. The design goal of the presented hardware is to create small and optimized computational units (SbS inference populations) that simulate an IP. While it is possible to realize a few of these SbS inference populations on a FPGA, the long term goal is to build ASICs (or networks of these ASICs) that allow to realize such a network where every IP is represented by its own non-shared SbS inference population.

In the following we investigated how the algorithm can be realized as an ASIC. We tested the resulting VHDL implementation with Xilinx Virtex 6 LX550T-1FF1759 FPGA on 4DSP (now abaco systems) FM680 cards. The long term goal is to use this design for an ASIC, thus all FPGA specialized circuits (e.g. hardware DSP cores) or other intellectual property cores were not used. The only exception to this rule is block RAM (BRAM).

## 2 MATERIALS AND METHODS

### 2.1 Design goals

There are multiple ways to realize the required circuits. For example, the number of components necessary for the arithmetic units can be minimized by using sequential processing as well as reusing the arithmetic units as much as possible. Another approach would be to design the circuits such that the calculations are done as fast as possible (measured in clock cycles) under the penalty of using much more components in the process. The design goal for the results of this study is leaning strongly in the latter direction by using pipeline designs for the arithmetic units. The reason for this decision lies in the low clock speeds. Using FPGAs or ASICs allows us to design a computational environment for these SbS networks that allow fast updates of the latent variable. For this task, this architecture allows better parallelization than commercial general purpose CPUs.

For implementing a flexible multi-layer Spike-by-Spike network on chip, the following main ingredients are required:

a) SbS inference population: Each such population consists of a population of neurons that are in a competition which is realized by normalization. A inference population receives spikes as indices *s* of sending populations and uses them to update internal states *h*(*i*) of each population member *i* and weights *p*(*s*|*i*) as well as produce outgoing spikes which reflect its internal latent variable *h*(*i*).

b) Input populations: Input into a SbS network is given by probability distributions *p*_*µ,g*_(*s*) represented by populations of neurons *s*, where *µ* denotes the actual pattern and *g* enumerates the input population. These probability distributions are used to generated spikes which are send to SbS inference populations for further processing. Input patterns can be e.g. pixel images, time series, or waveforms. The input pattern is interpreted as a vector of numbers, which is transformed into a probability distributions by normalizing it according the L1 norm. Every one of these normalized numbers is represented by an input neuron. The higher the value which is stored in the neuron, the higher is the probability that this neuron produces the next spike. The input pattern is provided from outside of the FPGA / ASIC from sensors (e.g. camera) or from data storage devices and programmed into the input neurons via a data bus. Typically the probability distribution stored in the input neurons doesn’t change for an externally defined number of spikes (until the computation of this input pattern is done, then it is usurped by a new probability distribution). The neurons in the input populations don’t react to any spikes produced in the SbS network. Thus they can be considered simplified versions of the inference population where only the spike generating part remained but without any weights, learning or updating of internal variables through spikes.

c) Network communication fabric: Spikes need to be exchanged between populations.

Focusing on the calculations that a SbS inference population needs to perform, a set of equations have to be realized: The basic equation realizes the update dynamic for the latent variables *h*(*i*) based on an incoming spike. In (Ernst et al., 2007) it was shown that only the identity (i.e. the index) *s*^*t*^ of the subsequently active neuron needs to be taken into account:

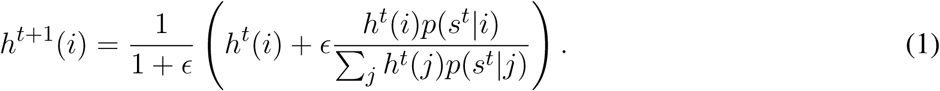

Besides updating the latent variables *h*(*i*), it is necessary to optimize suitable weights *p*(*s*|*i*) from training data for allowing the network to perform the desired function (e.g. pattern recognition). Two different approaches were found useful for learning weights. One is based on changing weights based on only single spikes observed during processing the actual pattern. This procedure is called online learning. The other approach utilizes information gathered over many spikes and several patterns, that is batch learning.

For online learning we focus on a multiplicative learning rule for the weights *p*(*s*|*i*):

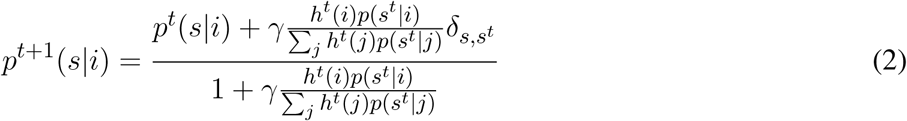

where *γ* is a learning parameter which can change during learning.

For batch learning, a variety of implementations exit, we focus on batch learning rules that base on

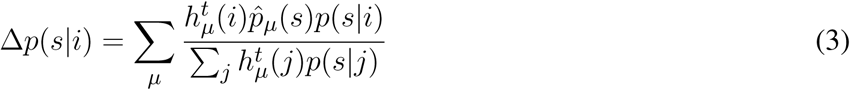

where *µ* identifies the training pattern, and where the sum may run over the whole or only parts of the complete set of training patterns. 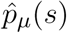 denotes the input probability distribution of the incoming spikes into the SbS inference population and is approximated by analyzing (counting) the spikes processed by the SbS inference population. For allowing a more flexible implementation of these type of rules, Δ*p*(*s*|*i*) is handed over by the FPGA to a CPU, which allows to implement batch learning rules based on

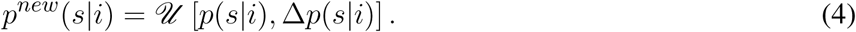

A simple example for an update rule falling into this category is

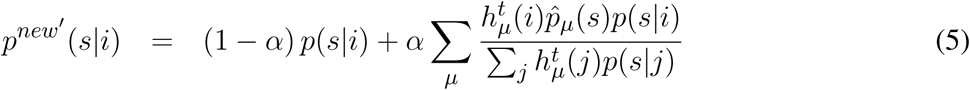

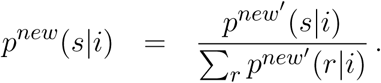

𝓊 could also be realized by Adam (Kingma and Ba, 2014) or L4 (Rolinek and Martius, 2018) using mini-batches. Since Δ*p*(*s*|*i*) is based on a multitude of patterns, offloading Δ*p*(*s*|*i*) to a separate CPU and performing the weight update on the CPU occurs at a lower rate than all other operations. Thus the reduction in performance by using a CPU for handling the weight updates is outweighed by the gain of flexibility.

Furthermore, we need the means to generate spikes from probability distributions or the latent variables using random numbers. Also specialized circuits are required for observing the spikes entering a SbS inference population and calculating the corresponding probability distribution 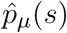 from these observations.

### 2.2 Non-negative numbers

Investigating the three main equations 1, 2 and 3 reveals that no subtractions are required and that no negative numbers appear. Furthermore most numbers (especially *h*(*i*) and *p*(*s*|*i*)) are in the range of [0, *…,* 1]. Incorporating these facts into the arithmetic units allows to simplify to the usual designs (Shirazi et al., 1995).

Taking the memory structure of the Xilinx Virtex 6 FPGA into account, where the block memory is organized into blocks of multiples of 1024 words with 18 bits each, we decided to used a custom variant of 36 bit floating point numbers as well as 18 bit fixed point numbers 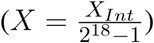.

In more detail, we designed the 36 bit floating point numbers as follows: Since only non-negative numbers are used, the usual sign bit was not necessary anymore. As part of batch learning, calculations on a typical CPU occur. This requires an easy way to convert our floating point numbers into their IEEE 747 counterpart. Thus we keep the number of bits for exponents, removed the sign bit and appended 5 extra bits to the lower significant bits of the mantissa. This allows conversion to be performed just by removing the not required bits of the mantissa or filling the extra bits of the mantissa up with zeros. We also introduced a similar derivative for 64 bit floating numbers (double precision) with 72 bits. Here 9 additional bits were added to the mantissa. These 72 bit floating point numbers are only used for representing Δ*p*(*s*|*i*) because summing over the contributions from larger amounts of patterns may otherwise lead to a degeneration of precision. For this reason, a similar extension from 18 bits to 36 bits was done for the fixed point numbers.

Figure S1 shows the coding for the two types of floating point numbers used in the design. The bits marked with gray boxes as well as the not available sign bit are different from the IEEE 747 standard. To simplify the floating point arithmetic units even more, only 0 as a sub-normalized number is allowed. In the supplemental materials, an investigation is presented where the impact of different representations on the performance of the MNIST SbS example network is analyzed.

### 2.3 Analyzing the equation’s structure

Analyzing the three equations 1, 2 and 3 reveals that all have common terms. Especially they share the term

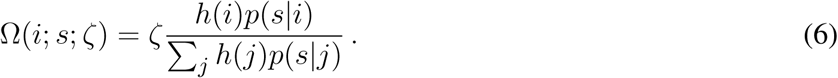

*ζ* can be ϵ, *γ* or 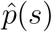. All three variants of *ζ* have in common that they change slower than *i*, which means that *i* can change many times before *ζ* is changed once. Also the *s* in *p*(*s*|*i*) changes slower than *i*. *s* can be selected through an observed spike *s*^*t*^ and only changes after all latent variables of that SbS inference population have been updated. Furthermore, for batch learning *s* should be selected as the slower changing index because this reduces the amount of times 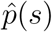 needs to be calculated.

Since memory is a scarce source on FPGAs and ASICs, it is advantageous to recall all the corresponding *h*(*i*) and *p*(*s*|*i*) pairs twice from memory during calculating Ω(*i*; *s*; *ζ*) while *s* is fixed. Since the arithmetic operation *Y/X* is known to be computational demanding, the following approach was chosen: 0.) All *h*(*i*) and *p*(*s*|*i*) pairs are streamed through a multiplication pipeline, thus making *h*(*i*) · *p*(*s*|*i*) available at the same speed with which they are recalled from the memory. However *h*(*i*) · *p*(*s*|*i*) is only available after a fixed delay.

1.) During the first sweep through the *h*(*i*) and *p*(*s*|*i*) pairs, Σ_*j*_ *h*(*j*)*p*(*s*|*j*) is calculated by observing the output from the multiplication pipeline. In the case of floating point numbers, adding two numbers take longer than two clock cycles. Figure S5a shows the steps necessary for adding these numbers and every step requires one clock cycle in our design. Thus we found that it is beneficial to implement the summation operation via an addition pipeline, where the output of the addition pipeline is feedback to its input (this procedure will be explained in detail later).

2.) After calculating 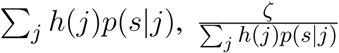 is calculated in a sequential fashion. This keeps the required amount of components for this arithmetic operation lower while a pipeline approach wouldn’t lead to a faster calculation anyhow.

3.) Finally, in a second sweep the *h*(*i*) and *p*(*s*|*i*) pair are recalled from memory. After the first multiplication pipeline which produces *h*(*i*) · *p*(*s*|*i*), a second multiplication pipeline is placed. The latter one multiplies 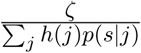 with *h*(*i*) · *p*(*s*|*i*) which results in Ω(*i*; *s*; *ζ*).

For reducing the amount of overall delay, it is beneficial to start step 3.) some time before step 2.) is fully complete. The timing has to selected such that 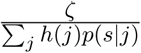 is just ready when the first *h*(*i*) · *p*(*s*|*i*) result leaves the first multiplication pipeline.

For the implementation of online learning, it is self-evident that the results from the circuits for step 0.) and 1.), calculated during the update of the latent variables, should be reused. Adding an additional sequential division unit and an additional multiplication pipeline allows to calculate step 2.) and 3.) fully in parallel to the ongoing update of the latent variables.

Since the calculation of Ω(*i*; *s*; *ζ*) of the batch learning update for the actually processed pattern isn’t done at the same time as the update of the latent variable, it is beneficial to use the same circuits for these two tasks.

## 3 RESULTS

### 3.1 Computational building blocks

Five basic building block can be identified for implementing the three equations. All five **Module**s have in common that input entering the **Module** is accompanied by an organizational index *i*. Delay lines in the **Module**s ensure that this index labels its corresponding output. Figure 2 shows an overview of the computations performed in these **Module**s. Table S1 lists the components required to map the circuits onto Xilinx Virtex 6 FPGA hardware. Resources listed are Slice Registers (687360 available on the used LX550T version), Look-Up-Tables (LUT, 343680 available), LUT Flip Flop pairs used (85920 available), and block Random-Access Memory (BRAM, 1264 blocks with 18k bits each or organized as 632 blocks with 36k bits). Every type of these resources represent highly complex circuitry (see Xilinx user guide UG364 ‘Virtex-6 FPGA Configurable Logic Block Table’ for more details). Every pipelined operation is designed such that one new operation can be started per clock cycle. However, it typically takes more than one clock cycle (also called latency) before the result of an operation leaves its pipeline. Furthermore some **Module**s contain sequential components that need a given number of clock cycles before they are initialized for their task and can perform their actual computation. Table S2 gives details concerning the latencies or how long the operations take.

**Figure 2.**
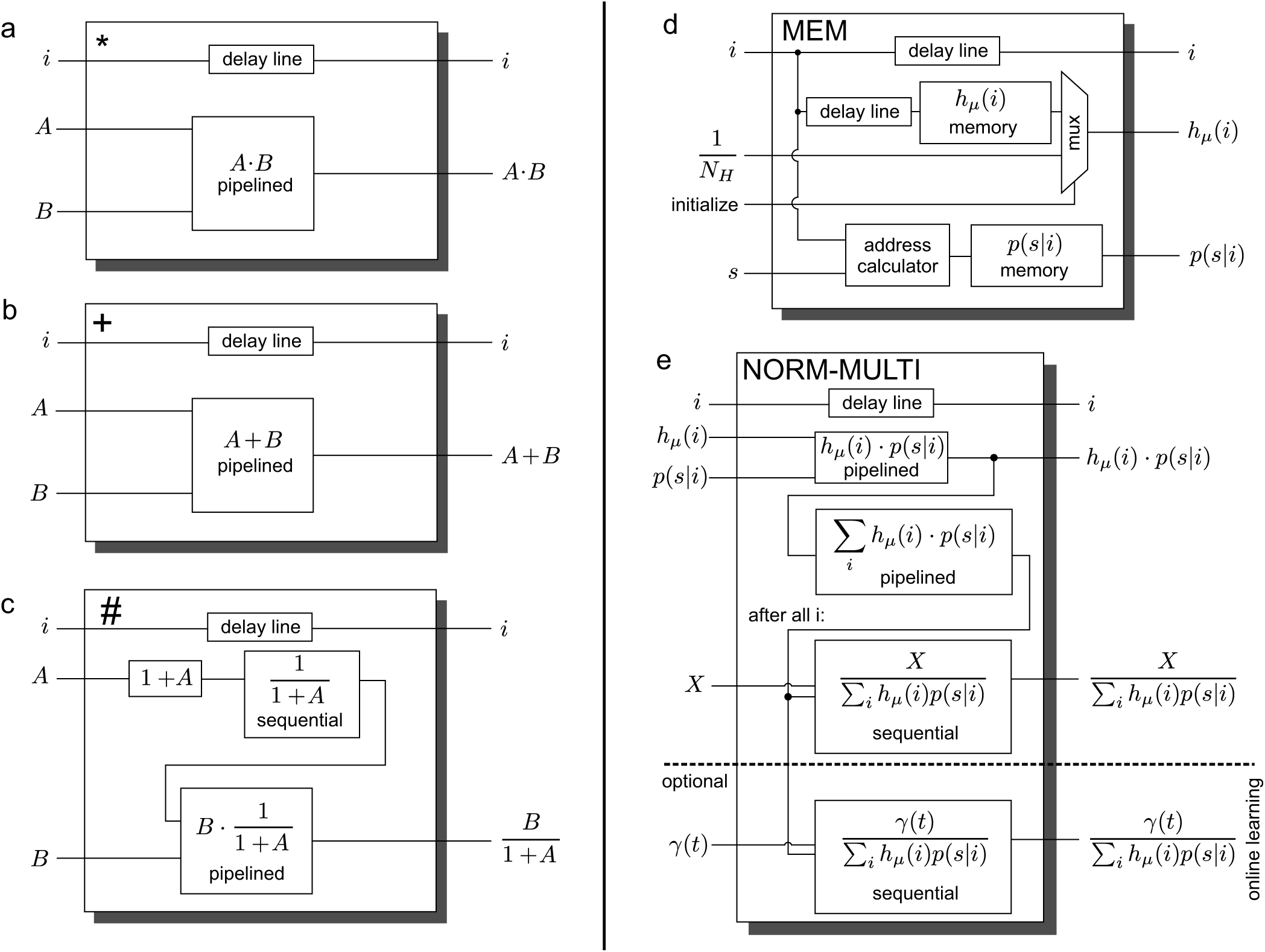
Basic computational building blocks: a.) **Module** *, b.) **Module** +, c.) **Module** #, d.) **Module MEM**, and e.) **Module NORM-MULTI**.

**Module** * (figure 2a) represents a simple multiplication pipeline. Two inputs *A* and *B* enter the Module and the result *A* ·*B* leaves the **Module**. In the case of fixed point numbers, the output has twice the number of bits compared to the input.

**Module** + (figure 2b) represents a simple adding pipeline. Two inputs *A* and *B* enter the **Module** and *A* + *B* leaves the **Module**. In the case of fixed point numbers, the output has one bit more than the input.

**Module** # (figure 2c) computes 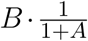, where *A* is assumed to be changing on a slower time scale than + *A* is calculated and then 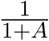 is determined in a sequential fashion. This result is used as one factor for a multiplication pipeline, while the fast changing *B* is the other factor. For fixed point numbers, the result of 1+*A* increases by one significant bit over *A* and the reciprocal operation doubles the number of bits again. The multiplication pipeline doubles the number of bits a second time, before the results exiting the **Module** is cut down to the original number of bits for *A*.

**Module MEM** (figure 2d) stores the latent variables *h*(*i*) and the corresponding weights *p*(*s*|*i*). The main components of this **Module** are two **block RAM Module**s (which can have different sizes), which provide a vector of memory values each. For each of the **block RAM Modules**, only one reading operation and one writing operation can be done in parallel during one clock cycle. While the access of the *h*(*i*) memory could be done directly, for *p*(*s*|*i*) the memory location needs to be calculated via an address calculator from *i* and *s* first (this is done according to the equation: Linear memory position(*s, i*) = *s* + *N*_*S*_ · *i* with *N*_*S*_ as the biggest possible *s* plus one if *s* and *i* are zero-based variables.). Thus the **Module** contains two address calculators, one for writing and one for reading operations. Since the calculation of the memory address takes some clock cycles (one multiplication and one addition) and the output of the **Module** are pairs of *h*(*i*) and *p*(*s*|*i*) based on the same *i*, read requests to *h*(*i*) memory are appropriately delayed. Figure 2d shows in a simplified fashion the reading part of the **Module**, for which a multiplexer is placed at the output of the *h*(*i*) memory. This multiplexer allows to usurp the *h*(*i*) memory output by an initialization value without writing it into *h*(*i*) memory first. For floating point numbers, at the input and output of the **Module**, the first bit of the mantissa is removed or re-added (see figure S1). This bit is necessary for calculations but is directly defined by the exponent and hence doesn’t need to be stored in memory.

**Module NORM-MULTI** (figure 2e) is a modification of **Module** *. In addition to the multiplication functionality, the cumulative sum over the output of the multiplication pipeline is calculated. In case of fixed point numbers, adding two numbers can be done in one clock cycle. However, for floating point numbers it takes several clock cycles for adding two numbers. In combination with receiving one new output from the multiplication pipeline in every clock cycle, this poses a problem. As a solution, we used an adder pipeline feedback on itself (see figure S5). The cumulative sum goes through three stages for floating point numbers: For stage 1 and 2, input *A* into the adder pipeline is defined by the output of the multiplier pipeline. In stage 1, which lasts as many clock cycles as required to pass through the adder pipeline, *B* are set to 0. Then stage 2 is entered where *B* is set to the output of the adder pipeline and thus creates a kind of circular buffer. After the last input value from the multiplication pipeline’s output was received (defined by an external constant), stage 3 is entered. In stage 3 *A* and *B* is set to 0 by default. The output of the adder pipeline is collected until two valid outputs are available. These two values are send as an *A*-*B*-pair through the adder pipeline. This is done several times and thus combines more and more remaining pairs until one value is left, which is the desired output of the cumulative sum. Finally, an external value (in figure 2e *X* is a placeholder for ϵ or 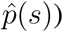 is divided by the actual cumulative sum in a sequential way. In the case of online learning the weights it is beneficial to perform a second division with the learning parameter *γ* in parallel. A note concerning the number of bits for the fixed point case: the cumulative sum cannot exceed 1 since *h*(*i*) is normalized to 1 and the values of the weights obey 0 ≤ *p*(*s*|*i*) ≤ 1. Since ϵ values which are larger than 1 can be interesting for some applications, the design allows ϵ values with up to 22bits (which allows for values up to 16). Thus 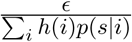 has 22+18 bits.

Besides the basic computation building blocks for calculating the three main equations, additional **Module**s are required for realizing the network. In particular a **Module** that allows to generate spikes from a probability distribution and a random number, a **Module** that analyzes spikes and calculates a rate 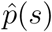 out of them, and a **Module** that offsets the weights in a normalized fashion during learning. The figures S2, S3 and S4 show simplified schematics for these **Module**s. Table S3 shows the amount of components necessary to build them as well as the number of clock cycles they require.

**Module Spike Generator** (figure S2a) converts a probability distribution and one random number into one spike. A spike is an index describing a position in the probability distribution that elicited the spike.

The values of the probability distribution (e.g. *h*(*i*) or normalized input pattern) are presented to the Module sequentially.

The Module sequentially calculates the cumulative sums over the observed part of the probability distribution and compares it to the random number. Is the actual value of the partial cumulative sum equals or is larger than the random number, the index of the probability distribution value that just contributed to the sum is the desired index. If every value of the probability distribution was shown but the last value and no index was found yet, then the wanted index is the last position in the probability distribution. In addition, for this design the values of the probability distribution are converted into a fixed point representation, since adding two fixed point numbers can be done in one clock cycle and the available random numbers are in a fixed point representation anyway.

**Module Spike Generator with offset** (figure S4a) Sometimes, especially during annealing while learning, it is helpful to add an offset *α* to a probability distribution and to fade out this offset over time. One way to generate spikes with an offset is to add the offset to the probability distribution, re-normalize it and then draw spikes from it. In the context of these circuit designs, a more efficient way is to rely on a double stochastic process using two random numbers. One random number is used to draw one spike from the original probability distribution without offset and one spike from a uniform probability distribution. The second random number is compared to *α*. The outcome of this comparison decides which one of the two spikes is used.

Random number generator (figure S3): For generating spikes, we need random numbers. For generating those, we use a Mersenne twister (MT 19937) which produces 32 bit random numbers (with a period of 2^19937^). A requirement for the MT 19937 implementation was that it needs to produce one random number in every clock cycle. Several different designs for such MT circuits are available. We decided to combine the designs from (Tian and Benkrid, 2009) and (Saraf and Bazargan, 2017). Our design uses two block RAM (624x 32bit words each) and 624bit logic RAM as memory. While it uses one more block RAM than such an implementation ultimately needs, it allows us to produce exactly the random number series in the order produced by Matlab or C++. Our **MT Modules** also contains a computational unit that calculates the initial table after reset given a seed value.

**Module Rate Calculator** (figure S2b) For batch learning the weights, the input rate 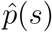 is necessary. For calculating the rate 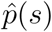, the following is required: The total number of spikes *c*_*All*_ observed and the number of spikes seen in each channel *s* counted as *c*(*s*). After an externally defined number of spikes has been counted, the rate is calculated and recalled by an external **Module**. This is realized by calculating 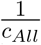 first and then multiplying it by *c*(*s*). Depending on the required representation of the rates, conversion into other number formats may be necessary.

**Module Normalized Weight Offset** (figure S2b) During learning it may be necessary to keep the weights *p*(*s*|*i*) away from 0 (otherwise the multiplicative learning algorithm may get stuck). Hence this Module adds an offset 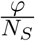 to each weight while they pass through this Module. However, the Module must also ensure that the normalization of the weights Σ_*s*_ *p(s|i)* = 1 is still kept intact. Thus the Module also divides the intermediate weight by 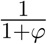 before it exits this Module. In addition, the corresponding *h*(*i*) values are delayed to keep them in sync with their *p*(*s*|*i*) counterparts. The **Module Normalized Weight Offset** is directly placed in series after the **MEM Module**. To keep the figures 3, 4, and 5 clear, we omit showing the **Normalized Weight Offset Module** and show only the **MEM Module** and not the combination of both Modules.

**Figure 3.**
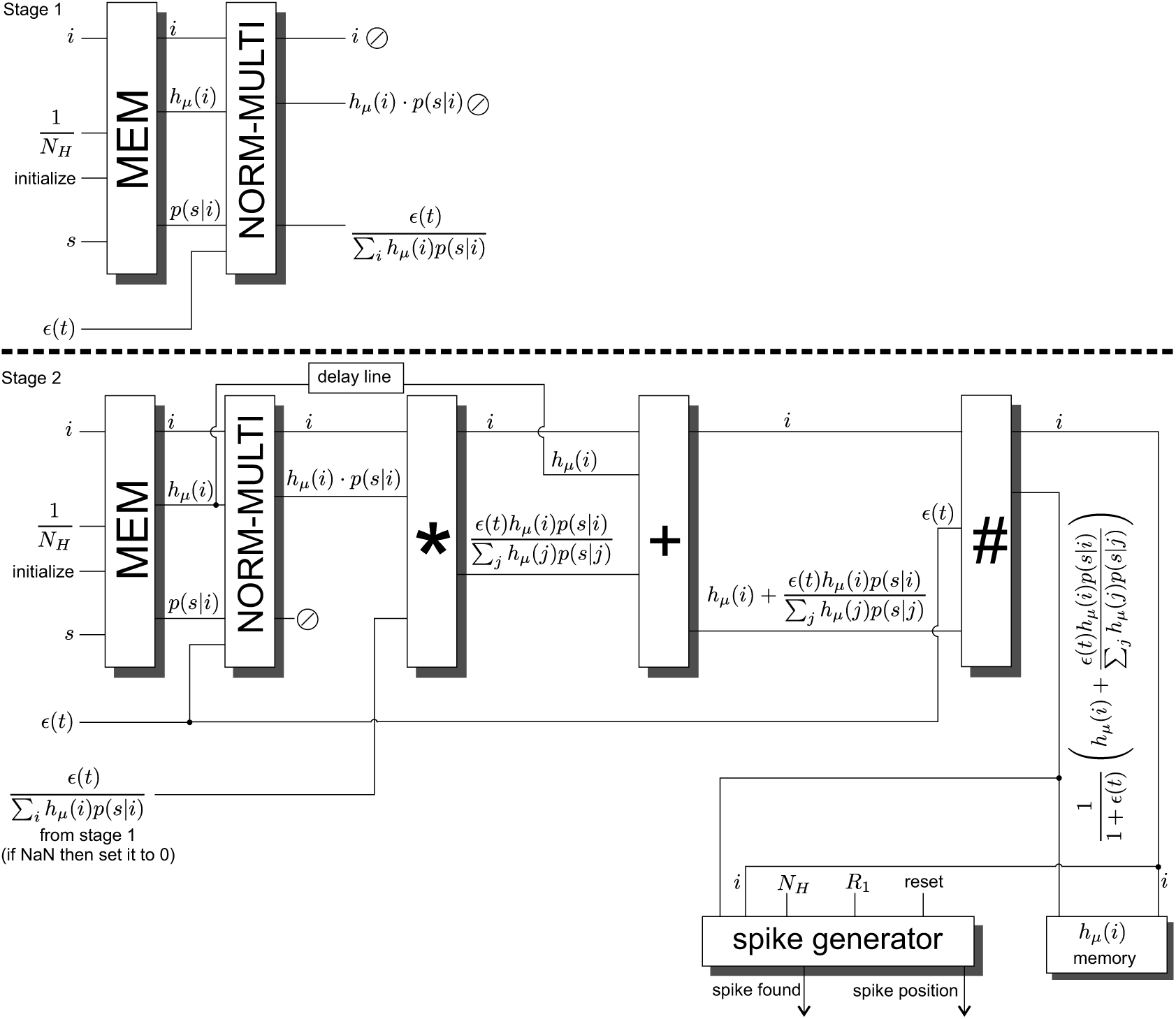
Circuits for updating *h*(*i*) based on a spike *s*. The update is done in two stage.

**Figure 4.**
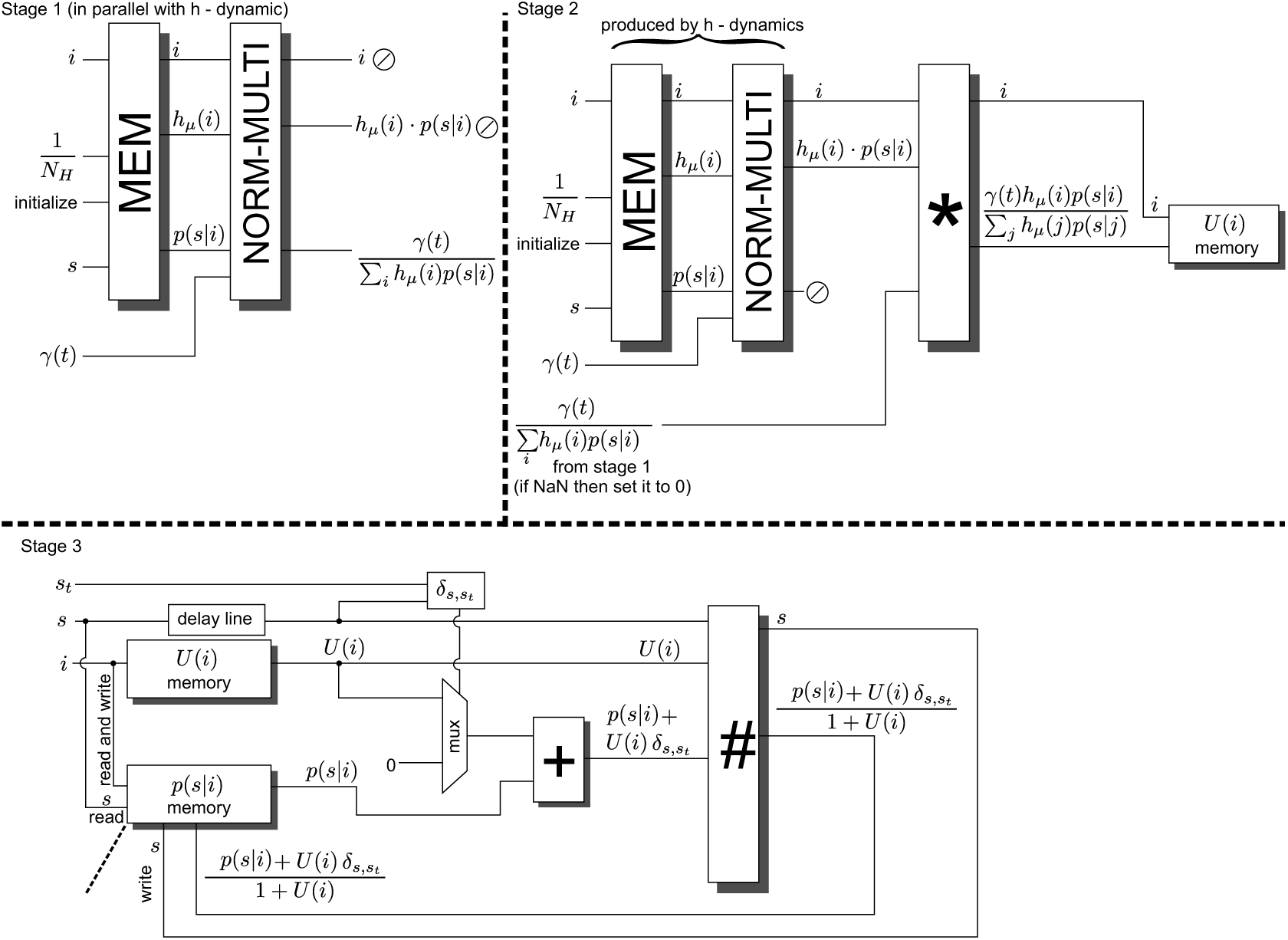
Circuits for updating the weights *p*(*s*|*i*) via online learning. The update of *p*(*s*|*i*) is done in a three stage process.

**Figure 5.**
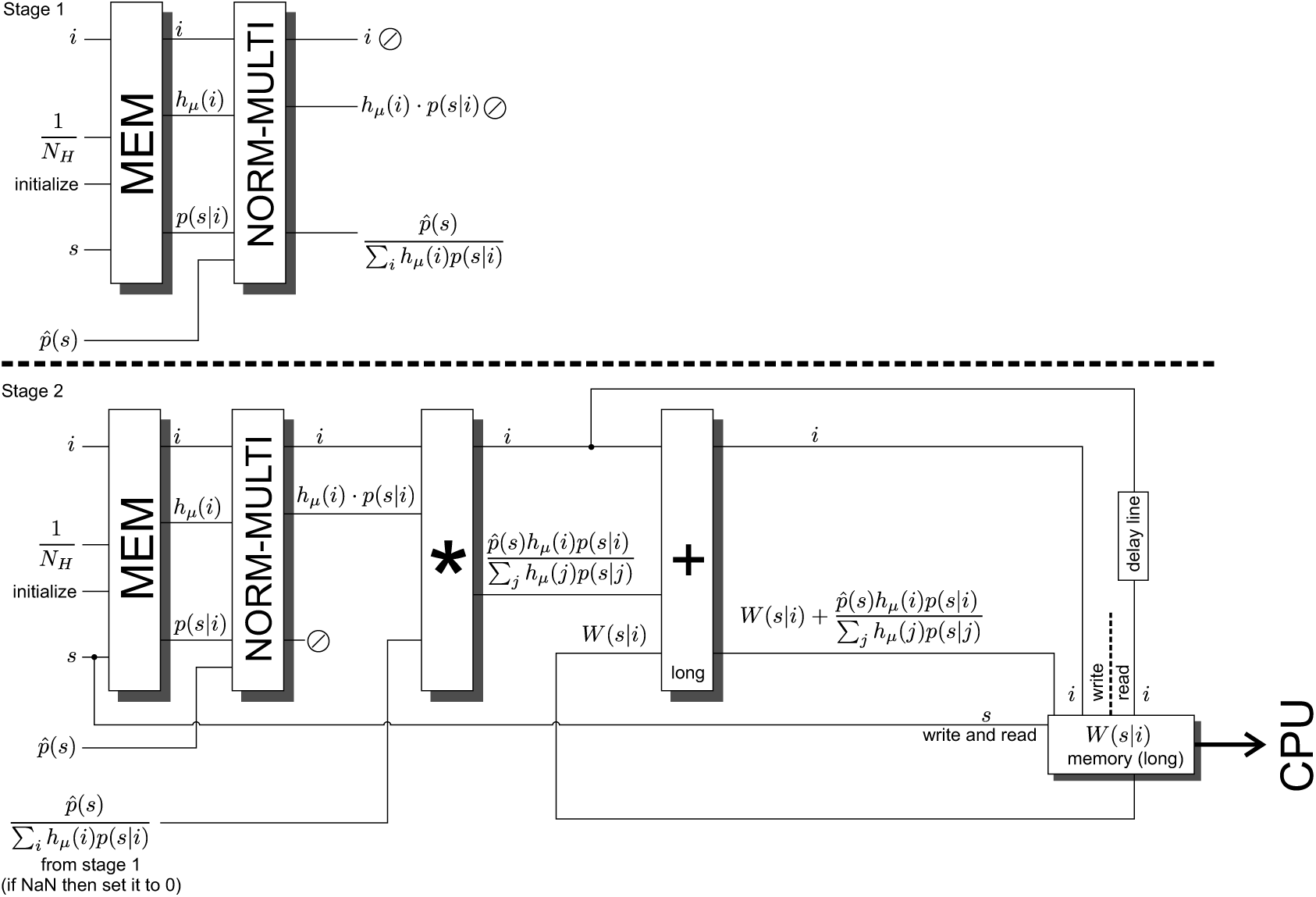
Circuits for batch learning. The matrix *W* (*s*|*i*) is calculated from many spikes and input patterns. The *W* (*s*|*i*) is send to an external CPU and used to calculate the new weights *p*(*s*|*i*). The calculation of *W* (*s*|*i*) is done in two stages.

### 3.2 Circuits for updating *h*(*i*), online and batch learning *p*(*s*|*i*)

With the presented building blocks it is now possible to implement the three equations 1, 2 and 3 as circuit diagrams. Concerning the update of *h*(*i*) based on an observed spike *s*, figure 3 shows how the building blocks are used. The update is done in two stages, during which the spike *s* is constant. In the first stage, the index *i* counts through all allowed values 1, *…, N*_*H*_. This recalls *N*_*H*_ pairs of *h*(*i*) and *p*(*s*|*i*) values. After an optional normalized offset is added to the weights *p*(*s*|*i*), these pairs a feed into a **NORM-MULTI Module**. As a result 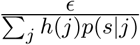 is calculated. Now stage two begins, with recalling the *h*(*i*) and *p*(*s*|*i*) a second time from the memory. In this stage, the output of the multiplier pipeline *h*(*i*) · *p*(*s*|*i*) of the **Module NORM-MULTI** is combined with the previous calculated 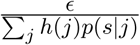 via another multiplication pipeline into 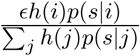. In turn, this intermediate result is added via an addition pipeline to its *h*(*i*) which was delayed to be available at the right moment in time. As a last processing step, the output of the addition pipeline is divided by 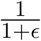 via a **Module** #. The output of the **Module** # dlivers the new *h*(*i*) values. These are then written into the *h*(*i*) memory of the **Module MEM**. Furthermore a **spike generation Module** (we used the ‘with offset’ variety in our design) observes the new *h*(*i*) values in parallel and, if provided with a random number, draws a spike out of the new *h*(*i*) distribution. Table S4 shows the number of components required to map this circuit onto the Xilinx Virtex 6 FPGA as well as how many clock cycles of latency the **Module**s *, + and # adds to the processing time.

Measured from the time then the first pair of *h*(*i*) and *p*(*s*|*i*) values are requested from **Module MEM** until the clock cycle when the last updated *h*(*i*) values is written into memory, the *h*(*i*) update takes 79 + 2 · *N*_H_ clock cycles for floating point numbers and 118 + 2 · *N*_H_ clock cycles for 18bit fixed point numbers.

In parallel to the *h*(*i*) update, an online update step for the weights *p*(*s*|*i*) can be performed. Figure 4 shows the three stage process for doing so. Stage one and two of the *h*(*i*)-update and online *p*(*s*|*i*)-update overlap. While 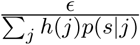 is calculated in state one on the *h*(*i*)-path, 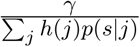 calculated in parallel. In stage two, the output *h*(*i*) · *p*(*s*|*i*) of the **Module NORM-MULTI** from the *h*(*i*)-path is multiplied via another multiplication pipeline with 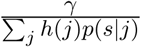 in parallel. This results in 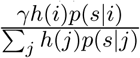 as an intermediate result which is stored in a temporary memory *U* (*i*). Furthermore the spike *s* is stored as *s*_*t*_ for stage three. In stage three the old weight values need to be updated with *U* (*i*) and stored as new weight values into their memory in **Module MEM**. For this process all *p*(*s*|*i*) have to be recalled from **Module MEM** (and optionally modified with an normalized offset via the corresponding **offset Module**) but with *i* as the slower changing index. For a given index *i* the **Module** # is prepared with the corresponding *U* (*i*). After this preparation is finished, the index *s* counts from 1 to *N*_*S*_. In a first step, it is checked if *s* equals *s*_*t*_. If this comparison is true, an addition pipeline adds *U* (*i*) to *p*(*s*|*i*). Otherwise 0 is added to *p*(*s*|*i*). The output of the addition pipeline is then divided by 1 + *U* (*i*) via the already prepared **Module** #. This result is written as the new *p*(*s*|*i*) values into their memory in **Module MEM**. The components required to realize state three and the temporary memory *U* (*i*) is listed in table S4.

Since stage one and two of the online *p*(*s*|*i*) update are done in parallel to the *h*(*i*)-update and with less complex calculations, the number of clock cycles required for stage one and two are defined by the *h*(*i*)-update. Concerning stage three, we measured *N*_*H*_ · (124 + *N*_*S*_) clock cycles for floating point numbers and *N*_*H*_ · (91 + *N*_*S*_) clock cycles for 18bit fixed point numbers. It needs to be noted that the reported clock cycle count for the *h*(*i*)-path and one *p*(*s*|*i*)-path were measured with the **normalized weight offset Module** in place. Furthermore, it is important to point out that the listed 124 and 91 clock cycles are mainly a result of the time during which the **Module** # needs to be prepared with the actual *U* (*i*). Using additional **Module**s # and switching between them would reduce these clock cycle counts significantly, however, would cost much more components.

Another approach to learn weights is to perform batch learning. In this learning mode, for a given input pattern *p*(*s*), *h*(*i*) is updated with many spikes *s*. Based on the final *h*(*i*) and the probability distribution *p*(*s*), for generating the observed spikes *s* in the first place, a contribution to the update of the weights from the actual pattern is calculated. Since *p*(*s*) is not available to this circuit, an estimate 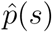 is calculated via the **Module Rate Calculator** from the observed spikes *s*. The circuit processes several patterns and accumulates these individual contributions into a matrix *W* (*s*|*i*). After the scheduled patterns are processed and their contributions are collected, *W* (*s*|*i*) is sent to an external CPU for updating the weights *p*(*s*|*i*). Using an external CPU for this purpose allows for a high flexibility on the realized update rule. Since the collected contributions can stem from a large number of patterns, *W* (*s*|*i*) is realized with twice the bits as *p*(*s*|*i*) for accommodating much larger numbers. The circuit for calculating *W* (*s*|*i*) in a two stage process is shown in figure 5. The required amount of components for stage two are listed in table S4.

For stage one of adding a pattern’s information to *W* (*s*|*i*), the circuitry of the *h*(*i*)-update can be reused, since there are no *h*(*i*)-update during this time. First, 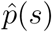 is calculated for one *s* by **Module Rate Calculator** and then used instead of ϵ. As result we get 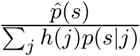. In stage two, analogous to stage two of the online *p*(*s*|*i*) update path, a multiplexer pipeline is used to create the intermediate results 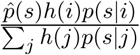 for all *N*_*H*_ indices *i*. The result of this multiplier pipeline is then converted into the corresponding number format with more bits and added to the content of the matrix *W* (*s*|*i*). After all *N*_*H*_ indices *i* are processed, *s* is incremented by one and the next 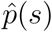 is recalled. For this new*s* the process starts at stage one again. This sequence of fetching 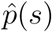, stage one and stage two is repeated until it went through all *N*_*S*_ index values of *s*. Altogether this takes 51 + *N*_*S*_ · (185 + 2 · *N*_*H*_) clock cycles for floating point numbers and 48 + *N*_*S*_ · (124 + 2 · *N*_*H*_) clock cycles for 18bit fixed point numbers.

After implementing these circuits on a FPGA, we investigated how the results differ from a Matlab simulation. Two factors contribute to the differences between the results for these circuits and a Matlab simulation: For all type of divisions, we neglected rounding the least significant bit of the mantissa of the result by just truncating it. Rounding would have required additional clock cycles. And for floating point numbers, the resolution of the mantissa is higher than the ‘single’ counterpart in Matlab simulations.

Four simple tests have been performed: a.) A non-normalized random *h*(*i*)-vector with *N*_*H*_ = 11 entries was normalized. b.) Given a random input distribution with *N*_*S*_ = 16 neurons produced 10 spikes. Using a given random weight matrix *p*(*s*|*i*), added with an offset through the **weight offset Module**, the latent variable *h*(*i*) with *N*_*H*_ = 11 was updated with these 10 spikes. c.) Using the setup of b.) after processing the sixth spike the weights *p*(*s*|*i*) were updated with every spike in an online fashion. d.) Using the setup of b.), after processing the 10th spike an update of *W* (*s*|*i*) was calculated.

For floating point numbers we found a maximum relative (difference divided by the Matlab simulation result) error around 3 · 10^−8^ for *h*(*i*), *p*(*s*|*i*) and *W* (*s*|*i*). For the 18bit floating point numbers, calculating the maximum absolute difference for *h*(*i*), *p*(*s*|*i*) and *W* (*s*|*i*) between the FPGA implementation and the corresponding Matlab simulation resulted in single digit differences, typically in the range from 0 to 6 for numbers in the range of [0, 2^18^ − 1]. These differences are a result of combining many differences created by neglecting rounding the results for divisions.

### 3.3 Connecting Spike-By-Spike inference populations and input populations

Typically, a network consists of more than one Spike-By-Spike inference population which need to exchange spikes. Or the task requires input patterns that need to be converted into spikes, which can be accomplished by an input population (a simple combination of block RAM for the input probability distribution and a **Module Spike Generation** for drawing spikes; see table S5 for a component count for a probability distribution of up to 1024 32bit values). As result a communication fabric between these spike producing and spike received network elements is required. Figure 6 shows the communication fabric that was implemented into the presented design.

**Figure 6.**
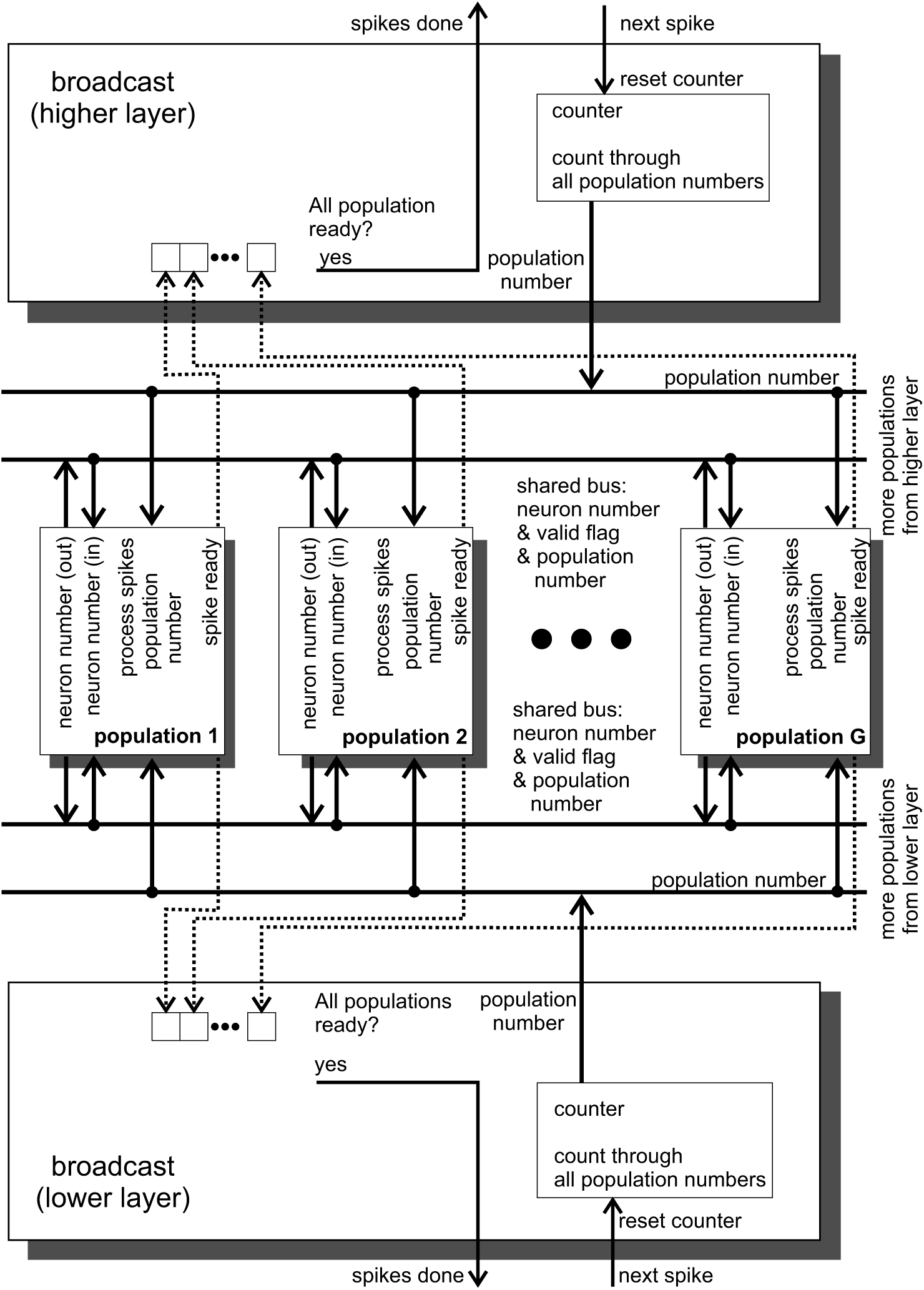
Communication between spike-processing elements in the network via broadcasts.

The communication fabric functions as follows: The input populations and Spike-By-Spike inference populations are connected to one or more shared data buses. The data bus conveys a unique identifier for the network element on that particular data bus, a neuron number for describing which neuron in that element produced a spike as well as a valid flag signaling that there is valid information on the bus at that moment. By default a network element is silent (all outputs are low). Every data bus owns a **broadcast Module** (see table S5 for a component count) which coordinates the activity on its data bus. When all **Module**s on the bus are ready for exchanging spikes, the **broadcast Module** sequentially calls all unique identifiers of the elements connected to this bus. Every element that sees its identifier and has a spike to report, sends out its own identifier and neuron identity that caused the actual spike. After the exchange is performed, the **broadcast Module** informs the connected network elements to process the information they received. The **broadcast Module** itself is controlled from a controller on a higher level. The shared data bus is realized by XOR elements which ensures that only one network element at the time uses the data bus. For some applications it is helpful to use more than one data bus for selected groups of network elements for keeping the required time to exchange spikes as low as possible. The presented design realizes two data buses.

Concerning the bandwidth of this spike communication, for every spike three numbers are exchanged: 1) The question if a spikes was produced by population *X* (where *X* is a 10 bit number plus one valid bit). 2) The answer of population *X* contains the 10 bit *X* and one valid bit itself as well as 3) the spike *s*^*t*^ which is a 10 bit number too, which is indexing the neuron that produced the spike in population *X*. However, population *X* only answers if it has a spike to report. The data bus is designed that transferring the question as well as the answer uses only one clock cycle. The questions and answers are transmitted non-blockingly in parallel on different parts of the bus. Assuming a population size of *N*_*H*_ = 1024, we can expect the FPGA to process ≈ 50,000 spikes per second and SbS inference population. For every spike, a total of 4 Byte per connected population or 200 kByte per second and connected population are exchanged.

While an input population doesn’t react to spikes on the shared data bus, a Spike-By-Spike inference population may need to update its internal variables according the observed spikes on the data bus. Figure 7 shows how incoming information on the data buses is handled by a Spike-By-Spike core. Assuming that it sees a valid set of information on the data bus, it compares the network element’s unique identifier with a dynamically programmable white list (table S5 details the required component for such a list). In case the unique identifier is on the list, the incoming information is processed further. Otherwise the incoming information is ignored. For white-listed information, the neuron number for the spike is stored and a flag that there was information received from this sending network is set. After the **broadcast Module** signals that all spikes have been exchanged and processing can begin (via a ‘process spikes’ line), the Spike-by-Spike core goes through its white-lists (one per data bus) and processes all flagged entries. It needs to be noted that the white list contains more information: An offset value that allows a dynamically programmed offset on the spike index *s* for this source of spikes as well as individual ϵ and *γ* values for every one of the white-listed spike sources.

**Figure 7.**
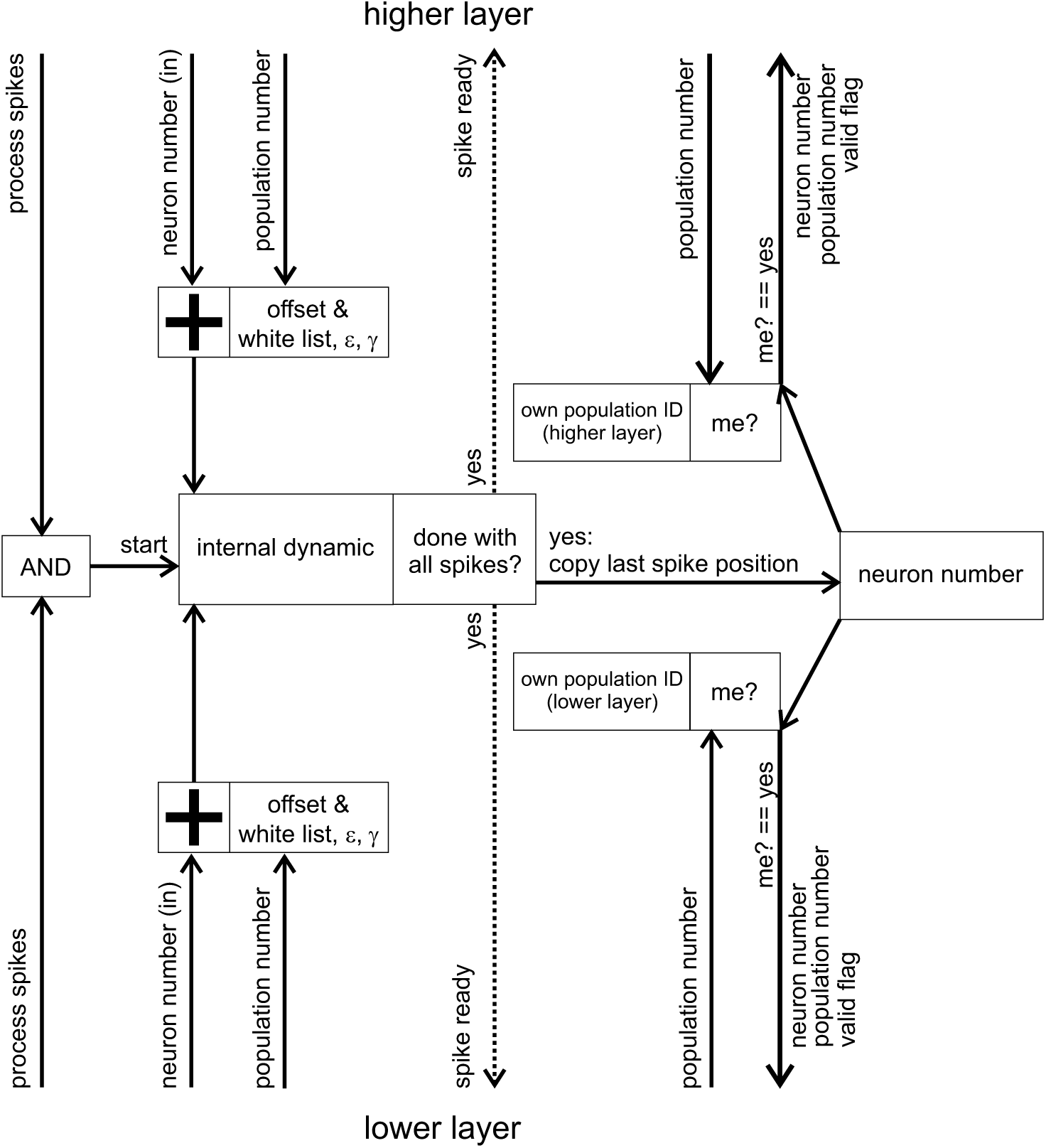
Handling incoming spike that enter a Spike-by-Spike inference population.

All the spikes in the white-list are processed which causes an update of the Spike-By-Spike inference population’s *h*(*i*). For every processed spike, automatically a new spike is drawn from the updated *h*(*i*) probability distribution. However, only the last one of these spikes is stored. After all spikes are processed a ‘spike ready’ line is raised, informing the **broadcasts Modules** that a new spike is ready for exchange. In the moment when the Spike-By-Spike inference population sees it’s own unique identifier on the shared bus, it puts its own identifier and the generated spike on the bus.

### 3.4 Coordinating the networks and exchanging data

While spikes are exchanged on specialized shared buses, also other data are required for operating the network (e.g. input pattern distributions *p*(*s*), weights *p*(*s*|*i*), white-lists and random numbers). Some data is generated inside the network, while other data needs to be provided from outside of the network. Furthermore, data needs to be read out of the variables from the network (especially *h*(*i*) distributions after processing input pattern as well as weight matrices *p*(*s*|*i*) or *W* (*s*|*i*) after learning) at externally defined moments in time. Thus we implemented two shared 32bit data buses: The first one for exchanging incoming external data as well as internally generated data (mainly random numbers and control sequences). The second one for sending data (e.g. results or information about the status of issued commands) to external receivers. See figure 8 for an overview.

**Figure 8.**
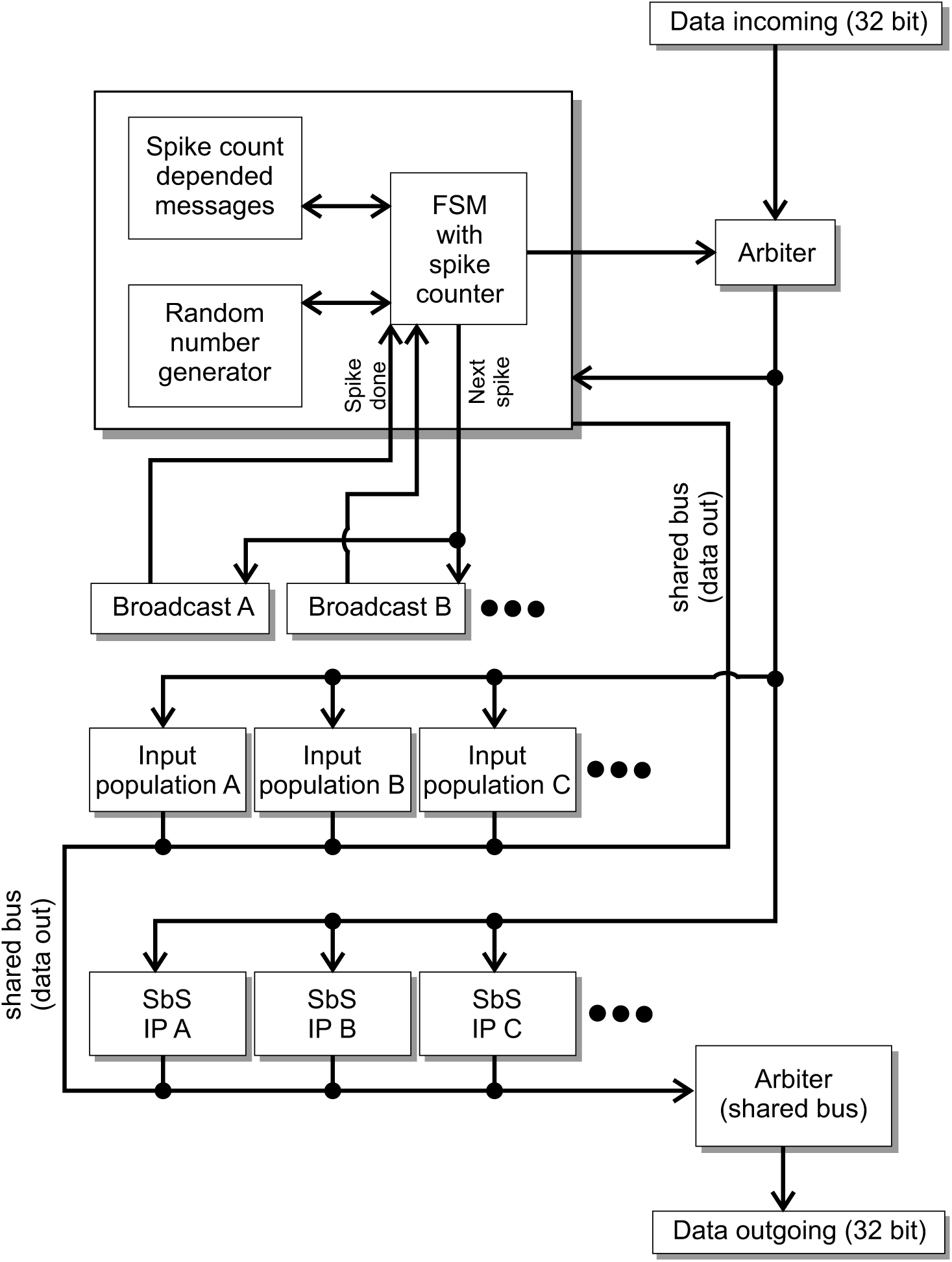
Data bus between the different elements as well as external participants.

This communication fabric was designed under the premise that information is loaded into the network (weight matrices, parameters and input patterns) and then the network runs for a number of given spikes (e.g. thousands of spikes). After that the results are readout from the network. New input pattern are loaded into the system, the network processes a number of spikes again, and the results are readout from the system again. It was not designed such that large amount of data is exchanged between every spike. Scalability of the communication fabric can be archived by splitting the communication fabric into parallel regions.

On the incoming data bus there are two sources for data: external data and internal data generated by a **message control center**. The **message control center** has several tasks: a.) Based on externally received information, it knows how many spikes the network needs to process as well as after which amount of spikes it needs to start with online learning of the weights. Hence it controls the activity of all the **broadcast Modules**. b.) It contains a Mersenne twister random number generator. After a spike, it provides all the network elements with new random numbers. It converts random numbers into messages and puts these messages on the data bus which programs the network elements with these random numbers. c.) It contains a mailbox system that is controlled by the number of already processed spikes. This allows to store every possible – otherwise externally provided – data control sequence in this mailbox together with a number of processed spikes that will release the message at that moment on the data bus. E.g. this message system can be used to change parameters like ϵ during processing the pre-defined number of spikes without waiting of any external just-in-time changes of parameters.

One arbiter joins the two data streams from the incoming data. Another arbiter combines all data sources that are destined for an external data receiver. In table S6 the amount of components required to realize all the participants on these data buses are listed. The network’s parts know 17 commands (e.g. set parameters, set *h*(*i*), *p*(*s*|*i*), read *h*(*i*), *p*(*s*|*i*), *W* (*s*|*i*), reset the **Rate Calculator Modules**, normalize *h*(*i*), set an input pattern probability distribution, set messages in the mailbox, and start the spike processing) that require up to nine 32bit words.

### 3.5 Test with a Xilinx Virtex 6 FPGA

We designed circuits for the network (see table 1 for details) such that it can be mapped, placed and routed without any special FPGA components (e.g. DSP cores), except for **block RAM Modules**, on a FPGA or ASIC. Our design, especially the number of buffering and pipeline stages, is optimized to generate a firmware that can be operated with 200+MHz clock speed on a Xilinx Virtex 6 LX550T-1FF1759 (speed grade: commercial 2). The Xilinx ISE 14.3 software is capable of producing such a firmware when the circuit is alone and the IO pins can be selected by the software.

**Table 1.**
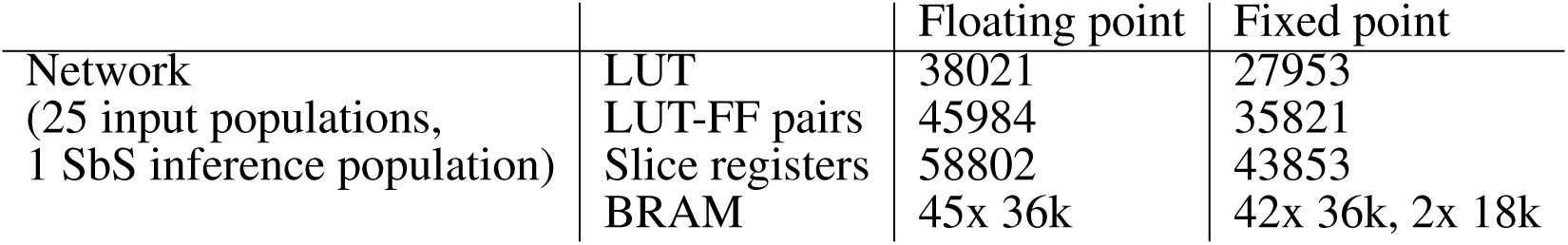
Amount of components required for a network with one Spike-By-Spike inference population and 25 input populations.

However, we weren’t able to achieve these high clock rate when the network was embedded into the third party firmware which is required for communicating with the Virtex 6 FPGA on the 4DSP FM680 PCIe card. We used the ‘adder’ example from the training materials and simply replaced the ‘adder’ core with our network. We allowed for a separate clock domain for the network by insulating the data-flow by dual clock FIFOs on the input and output side. Nevertheless, Xilinx ISE wasn’t able to provide us with a working firmware if we set the clock speed for the network to 200MHz. Thus we were forced to use the overall 125MHz clock provided by the 4DSPs example design also for our network.

With the 125MHz clock speed were were able to run the network on 4DSP FM680 cards under Linux (Centos 7.5 64bit with driver version 04.05.2018). Results were identically to the Xilinx ISim simulations. However, the overall design of the 4DSP FM680 doesn’t fit for our application very well. This type of card is designed for high data bandwidth applications where 64bit words in blocks of 1024bits are continuously exchanged. On one side we were forced to incooperate our 32bits into block of 1024bits. On the other side we had problems receiving data from the card because our network doesn’t produce data continuously. Thus the Linux driver and our software had to recover from trying to read from the card when it had no data for us. All this combined, slows down the communication with our network on the FPGA significantly.

In the supplemental materials we examine how the presented design fairs in comparison to Intel CPUs. In addition, it is investigated how the performance in the SbS MNIST example network (see figure 1) reacts to floating point and fixed point numbers with different number of bits. The conclusion of these tests is that FXP IPs are an interesting option, especially for IPs with smaller number of neurons. However, it needs to be tested if, in the given use-case, additional problems haven’t been created by the use of FXP IPs.

## 4 DISCUSSION AND CONCLUSION

The paper presents a design and an investigation of a circuitry optimized for implementing neuronal networks based on Spike-By-Spike (SbS) neurons (Ernst et al., 2007). As in the brain, signaling is based exclusively on spikes, interactions among local neuronal circuits are stricly non-negative (Dale’s law), it allows for recurrent interactions in cyclic architectures, and both, dynamics of the neuronal state variables as well as the learning rules can be local (Rotermund and Pawelzik, 2019b). Thereby this approach realizes a compromise between artificial neural networks and biologically realistic models employing detailed models of spiking neurons.

In a typical spiking neuron models (Izhikevich, 2004) are simulated with fine temporal resolution. In particular, simulations of networks of *noisy* leaky integrate-and-fire (IaF) neurons require representation of real time. Typically, the membrane potential needs to become updated every time step *dt* which is often in the range of sub-milliseconds. The number of updates between two spikes depends on the firing rate of that neuron. If for example *dt* = 0.1*ms* and the firing rate is 10Hz then this would translate roughly into 1000 updates of the membrane potential. In contrast, the SbS-approach avoids simulations of real time dynamics and would perform one update of a whole population between two input spikes. While the different types of spiking neuron models (Izhikevich, 2004) have varying number of computations for one update, in a SbS population with *N* neurons 3*N* multiplications, 2*N* summations, and one division are used for one update of the whole population. This reduction in computational requirement is payed by a decrease in biological realism.

In a SbS network time is only progressed with each spike received by an inference population (IP). Thus no computations have to be performed in between spikes, which drastically reduces the computational demand. An approach akin to the SbS’s removal of real time is also known for integrate-and-fire neurons. This so called event-based neuronal networks (e.g. Brette (2006, 2007); Serrano-Gotarredona et al. (2015); Lagorce et al. (2015)) use analytic solutions of the neuron’s dynamics to bridge the time between to spikes. However, with stochastic neurons the event-based approach becomes problematic (Brette, 2007). This is similar to the problem of finding an analytic solution for the first passage time (Burkitt, 2006a,b) for neurons with stochastic inputs in a network, which is a hard problem. For SbS networks, the stochasticity rather is feature because it corresponds to importance sampling of the input as well as the latent variables. This acts as a filter for capturing the more dominant information in the network and suppress noise.

In SbS networks, neurons are organized in populations where the neurons within a population compete with each other. Every neuron in a populations has a latent variable *h*(*i*). The value of *h*(*i*) is a positive number and the competition is expressed by a normalization over all latent variables of a population (Σ *h*(*i*) = 1). In between populations, neurons communicate exclusively via spikes (i.e. an index *s*_*t*_ describing a single neuron’s identity in a population at a time *t*). The connections between neurons from different populations are described by weights *p*(*s*|*i*) (with *s* as the emitting neuron and *i* as the receiving neuron). Also the values of the weights are positive numbers and limited in size by a second normalization condition (Σ_*s*_ *p*(*s*|*i*) = 1). Thus everything describing the state of the network is given by positive numbers in the range of 0 to 1. In this paper we compared how different types of representations for these numbers result in different sizes of computational circuits as well as different required numbers of clock cycles. In particular we examined 18 bit positive fixed point numbers and 36 bit positive floating point numbers. The latter are a custom derivation of the IEEE 754 standard by assuming positivity and a higher bit count for the mantissa. The fixed point representation requires less components but it was found that SbS inference populations can cause problems during learning, especially with larger numbers of neurons.

If a spike *s*^*t*^ is received by a population of neurons, for every neuron in this population an update of it’s latent variables is calculated through

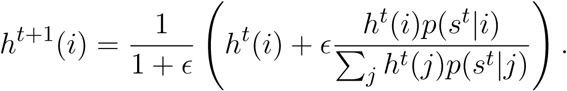

We presented computational circuitry for this equation that only requires to read the involved *h*(*i*) and *p*(*s*^*t*^|*i*) value pairs twice during the update process. For floating point numbers a number of 179 + 2 · *N*_*H*_ clock cycles and 118 + 2 · *N*_*H*_ clock cycles for the fixed point numbers were measured. The constant numbers of clock cycles in these two equations don’t scale with the number of neurons because they represent the time which the information needs to travel through the pipeline structure of the computational circuitry as well as setting up the circuits for incoming data. These amounts of clock cycles can be reduced if optional features are removed, like e.g. adding an normalized offset to the weights. Auxiliary circuits were designed such that, given a random number, they draw a new spike from *h*(*i*) while simultaneously updating the h-values. Furthermore it was shown that circuits for learning the weights (see equation 2 and 3) can be designed, while keeping the amount of required components low by partially reusing circuitry for the h-update.

Also for batch learning, circuits and memory are included into the presented design. Thus during batch learning, the contributions from many training patterns are collected within the circuitry. In the design phase of the system we decided against a hardware implementation of a specific learning rule that uses this accumulated data and calculates updated weights from it. Instead the idea is to use a CPU for these final weight update calculations, to allow large flexibility in choosing the ‘right’ variation of the learning rule later and allow for changes in the learning rule if new learning methods arise in the community without changing the hardware design. Combining FPGAs with CPUs has a long tradition, e.g. Intel started to offer 2010 the Stellarton (Series E6×5C, Atom CPU with FPGA). The Xilinx Virtex-4 FPGA from 2004 had versions with integrated PowerPC cores. At the same time, a soft core for the low-cost Xilinx Spartan-3 FPGA (a 32 bit RISC processor called MicroBlaze) was released as intellectual property core. This is also an approach which the neuromorphic community also adapted, e.g. (Wunderlich et al., 2019; Naylor et al., 2013).

Given a collection of populations of SbS neurons, the exchange between populations of spikes requires organization. In our framework we show that exchanging spike information can easily be done by a common data bus. A controller checks if all spike generating elements on that bus are ready for exchanging spikes and then calls every element on that bus to report its spike. Every SbS inference population has a programmable list which defines the spike producing elements it listens to. This allows user-defined changes of the network architecture on the fly. Also it allows to remove broken elements (which might be e.g. a result of production failures) from the network and to reallocate – if available – other resources to take its place. The lists contains additional information to allow for changes of the ϵ parameter for the h-dynamic and the *γ* parameter for online learning depending on the source of the spike. This feature is essential for more complex networks. In the same spirit, we introduced a ‘mailbox system’ into the framework. It allows to make processing a given pattern for a user-defined number of spikes independent from external sources. Sometimes it may be necessary to change the ϵ parameter after a given number of spikes. Without the mailbox system, an external controller would need to determine the state of the network (i.e. are the required number of spikes already processed?), then change the parameters, and tell the network to continue for another amount of spikes. This would obviously waste time. In comparison, with the mailbox system the network distributes the pre-defined messages about the parameter change automatically in the moment a pre-defined number of spikes have been processed. Obviously, the same is true for the required random numbers for producing spikes. Thus it is also essential that random numbers are sent to the consumers of these random number with as small as possible pauses. For this reason it was necessary to add a random number generator (Mersenne Twister 19937) to the overall message controller with its mailbox system. The message controller also takes care that the correct number of spikes is processed and that any external system is informed that the network finished the pre-programmed number of spikes.

As a second type of spike producing element in the network, we designed input populations. These input populations take a probability distribution, where every value is represented by a 32 bit fixed point number, as well as 32 bit random numbers and produces spikes from this information. For programming these probability distributions as well as configuring any other parameter or variable in the whole framework (or network) two data buses (one for incoming and one for outgoing data) with a pre-defined set of commands are introduced into the design. The aforementioned message controller receives commands from the incoming data bus but can also introduce its own commands into this data stream that is seen by all the elements of the network.

The presented design was transferred into a FPGA firmware written in VHDL. It was taken care that no special FPGA vendor proprietary Modules (except block RAM) were used, for allowing re-using the design for developing a custom ASIC. For proof of feasibility, we integrated the VHDL code into an example firmware for the 4DSP FM680 cards which hosts a Xilinx Virtex 6 FPGA from 2009. We successfully generated a binary firmware bit file for this card, tested it under Linux and compared it with the results from Matlab simulations. Small differences in our simple example simulation were found, which were expected due to the difference in precision (floating point numbers: 5 more bits representing the mantissa for the FPGA implementation compared to Matlab) as well as a different handling of rounding during the mathematical operation of division. We decided that the differences through rounding are not relevant, especially when compared to the required increase in the amount of components and processing stages necessary for reaching a perfect match with the Matlab simulations. While the pure SbS network would have been able to run with over 200MHz on an otherwise empty Virtex 6, combining it with the existing card manufacturer’s firmware unexpectedly forced us to reduce the clock speed down to 125MHz. Isolating the network into a separate 200MHz clock domain didn’t work out. Since the VHDL code was optimized for a 200MHz stand-alone implementation on a Virtex 6 LX550T FPGA, several buffering steps in the pipeline architecture were added which are unnecessary for a 125MHz operation. A second SbS inference population was added to the firmware’s network and tested out fine. Extrapolating from the number of required components for one SbS inference population, we expected that we could increase the number of SbS inference populations to 7 or 8 for floating point number on this Xilinx Virtex 6 LX550T. However, the generation of these firmwares failed in the Xilinx ISE design tool. Our guess is that the reason lies in the fixed placement of the some components (especially block RAM) on the FPGA. Increasing the number of SbS inference populations produces distances between components that are too long for even 125MHz on this FPGA from 2009. We didn’t pursued the increase in the number of SbS inference populations any further, because we found that the 4DSP firmware, driver and software framework for this card is optimized for continuously streaming of large bandwidth of data. However, in our application the data is send in a stop-and-go fashion which doesn’t ensure that data is always available. Thus it was necessary to probe the cards for available data via blindly trying to read data from the card and to rely on slow timeouts of the Linux driver if there was no data available at that exact moment. Furthermore we had to fill up our data streams with zeros (up to 31x bit in zeros for filling up compared to the payload) to conform to the required data format. In the end we decided to focus on transferring the design onto a custom ASIC instead of optimizing the FPGA firmware.

The computationally attractive aspect of a SbS neuron based network is the property that populations of neurons only communicate via spikes. Thereby every population of neurons operates independently from all the other populations. The internal computation of any given population is based exclusively on the incoming spikes and its internal variables. After finishing its computations every population of neurons also only sends out spikes as a signal to other populations. This locality property is true for updating the *h*(*i*) values as well as for learning the weights *p*(*s*|*i*). Thus the locality of its populations is the key ingredient for massively parallelizing networks with such elements (populations). The optimal way for parallelizing such networks is to provide independent memory and computational circuitry for every individual population of neurons. With the specialized processors proposed in this paper one update cycle in a large network of such populations does not scale significantly with the number of populations. This stands in stark contrast to running such networks on clusters of computers where one would experience latencies and transmission delays that scale much worse with the number of nodes due to the creation of network packages, their transmission and their analysis by soft-and hardware.

In future we will design a custom ASIC for networks with larger numbers of SbS inference populations. The desire for using ASICs compared to continuing using FPGAs stems from several reasons: The goal is to realize as many SbS inference populations as possible and give every SbS population in a network its own SbS inference population. However, this might require a network of ASICs. The main limiting factor with FPGAs is the amount of available memory (block RAM) and the corresponding routing problems. Every SbS inference population needs its own dual port memory, at least for its latent variables. On a FPGA this kind memory comes in block RAM Modules and these are fixed resources which are at certain manufacturer defined positions. These RAM blocks are spread over the whole FPGA die. If a SbS inference population needs more memory than one of these RAM Modules can provide, several of these RAM Modules are connected. This is a big problem for routing and timing which can be strongly experienced in our Virtex 6 FPGA design. It is a major factor in limiting the clock speed if several SbS inference populations are realized on the FPGA. The required logical components and the block RAMs are still available but the timing is too problematic due to the long distances. Furthermore, FPGAs are general purpose chips and the SbS inference populations have different requirements: The ration of logic circuitry to block RAM seems not to fit into the exception for general purpose case. Using ASICs allows us to place what we want in such a way that everything that needs to interact in a fast way is in close proximity (i.e. everything that is part of one SbS inference population) and everything that communicates over the spike bus can be placed further apart (and this communication can be on a much slower clock rate).

As a preparation we made sure that the design doesn’t use any special 3rd-party intellectual property cores. For improving the processing speed of the SbS inference population, it might be interesting for a custom ASIC to use different clock domains for exchanging spike information between populations and doing the calculations inside the SbS inference population. This would also ensure that the SbS inference populations can retain high computational speeds even if the distances between the populations on the chips for exchanging spike information increase. This even allows to extend the data bus for exchanging the spikes over a multitude of chips. Furthermore, it is also not necessary that all SbS inference population run at the same speed. It is enough that only the part of a population responsible for communicating with the data bus for exchanging spike information is in sync.

The area on chip required for memory may be an issue for the custom ASIC. Concerning the weights, in a convolution setting all populations in a layer use the same weight values. Thus in such an application it could be suitable to use a common RAM for all populations in that layer. The requirement with shared convolutional weights is as follows: For every update of the latent variables *h*, every SbS inference population gets a spike *s*^*t*^. For performing the required computations for the update, the weight vector *p*(*s*^*t*^|*i*) for all *i* ∈ [1, *…, N*_*H*_] is required. For example in the MNIST example, the convolutional weight matrix between the input and the first hidden layer has the size *N*_*S*_ = 50 × *N*_*H*_ = 32. Thus *p*(*s*^*t*^|*i*) is a vector with 32 numbers. However, there are only 50 different versions of this 32 number vector. On the other side, we have 576 individual SbS inference populations in the hidden layer 1, hence, 576 different spikes to process in every time step of the simulation with only 50 allowed values. The idea for the shared RAM is that all 50 vectors are sequentially recalled from RAM and broadcast to the 576 SbS inference populations. Every one of these 576 SbS inference populations know their own spike *s*^*t*^ and filter their 32 number vector from this common data stream. Obviously it takes 50x longer to do this broadcast procedure compared to the case where every SbS inference population has its own weight matrix. However, this reduction in speed in traded in for reducing the requirement for weight memory by a factor of 576. Furthermore, this would allow to use faster external memory Modules which would allow a broader broadcast bus (e.g. two or more vectors are broadcast in parallel). Or using custom RAM that delivers the *p*(*s*|*i*) values for all *s* during one read cycle in parallel would be beneficial, which would create *N*_*S*_ streams of weight values in parallel. In this scenario, every SbS inference population would have a multiplexer connected to the output stream of the RAM and would, according to the value of *s*^*t*^, switch between the different weight value streams.

The performance we can achieve in the MNIST benchmark (Rotermund and Pawelzik, 2019a) is 99.3% classification correct by using an error-back-propagation & momentum based learning rule developed for SbS networks which takes the requirements non-negativity and normalization into account. The learning rate was modulated by the expected remaining error estimated from the training data set. In addition, the fully connected layer H5 was subjected to a drop-out variant during learning. This learning rule is compatible with the presented hardware design. Using the same network architecture for a non-spiking neural network (Rotermund and Pawelzik, 2019a) we found 99.2%. These performance values are also reported to be comparable with traditional non-spiking convolutional neuronal networks (Tavanaei et al., 2018).

Using the presented SbS MNIST network with a local learning rule which is directly based on the gradient – like it is used in the supplemental materials – we still yield a performance of 97.3%. This is roughly the same performance we got using a classical non-spiking convolutional neuronal networks using the same network structure and as well with only the gradient (without any additional optimizers) (Rotermund and Pawelzik, 2019a).

Comparing our performance values with other networks (e.g. see for a list of MNIST networks (Tavanaei et al., 2018) and http://yann.lecun.com/exdb/mnist) is not as simple as comparing the values. In our case we didn’t optimized our network structure for the use of SbS inference populations. We rather choose to re-use the network structure associated with a Tensor Flow Tutorial because this gave us a base-line for a network design which our computer cluster was just been able to simulate. Furthermore, we didn’t used any input distortion methods (e.g. shifting, scaling, or rotating the input pictures) for increasing the size of the training data set. The reason was that this would have been to much for our computer cluster, like it would have been to optimize the parameters used in the SbS MNIST network. Or in other words: The performances shown for the MNIST SbS network doesn’t reflect what a fully optimized SbS network might be capable to deliver. Typically the performance values shown for neuronal networks are for an fully optimized network, like performances of 99.8% (Wan et al., 2013).

Due to the communication problem with the FPGA cards and the limited number of FPGA cards we have available (we own two of those cards and their price was 16,000+ Euros each), we had to perform the analysis of the MNIST network on a cluster of computer with Intel CPUs. It took our cluster 19 days with shy of 400 cores to perform the presented MNIST simulations. On the two FPGA cards with one SbS IP each, this would have taken approximately over a decade. This shows that the number of parallel cores is key in operating SbS networks in a fast fashion.

Obviously this raises the question why we are not just continue to use normal CPUs or migrate to GPUs instead of developing a special ASIC for that task. The two main aspects of the answer are parallelization (i.e. as many cores as possible) and memory bottleneck.

In the case of CPUs, the results of the timing measurements showed that as long as the required memory stays within the CPU’s L1/L2/L3 cache, the multitude of available cores can be efficiently used to simulating SbS IPs. The moment when the amount of memory required exceeds the CPU’s cache, the situation starts to change. Now the memory bandwidth to the external memory Modules starts to limit the SbS performance and the effective number of cores usable in every point of time dwindles. Thus even if a large number of cores is available on a CPU, only a few cores can be supplied with the required information continuously while the rest of the cores wait. This problem is amplified if one core is responsible for simulating several SbS IPs. In this use-case the information for the different SbS IPs (latent variables and weight matrices) need to be switched and loaded into the CPU’s cache continuously. Thus the memory bandwidth becomes the limiting factor again. In summary, CPUs are an excellent solution to simulate SbS IPs as long as the number of SbS IPs don’t exceed the core count of the CPU and as long as the required data fits into the CPU’s cache. For SbS IPs with large number of neurons and/or large weights matrices CPUs are especially a good choice because of the integration of CPU caches with several dozens of MByte.

There are special multi-core CPU ASICs with large core counts available. Examples are the Epiphany-IV (Olofsson et al., 2014) 64 RISC cores with 2 MB on-chip distributed memory (presumably 32 kByte per core) or the Epiphany-V (Olofsson, 2016) 1024 RISC cores with 64 MB on-chip distributed memory (presumably 64 kByte per core). As long as the latent variables and the corresponding weight matrix fit into the core’s local memory such an approach can be an interesting alternative for simulating SbS IPs. However, these types of ASICs are optimized for their amount of cores but were not designed to access external memory with high memory bandwidth. Thus, these multi-core CPU ASICs are a good alternative as long as everything necessary can be stored within the core’s memory. In the case of 32 bit floating point numbers this allows to store only one or two thousand values per core. Assuming 128 neurons in an IP, this would allow to connect every one of these neurons to only 15 input neurons. This back-on-the-envelope calculation excluded all the required auxiliary memory and space for program code necessary for an efficient computation which needs to be deducted from the available storage for the weights and latent variables too. However, the next generation of CPU from AMD are expected to contain core counts in the region of the Epiphany-IV but with much more on-chip memory.

Modern GPUs on the other hand are optimized for the use in memory and computational demanding applications, like e.g. deep non-spiking networks. The nvidia TU 102 GPU (see nvidia turing GPU architeture whitepaper WP-09183-001 v01 on http://www.nvidia.com) contains 72 (or 68 when on a RTX 2080 Ti graphics card) streaming multi-processors (SM) with 96 KByte of L1 cache / shared memory each. These 96 KByte can be configured as 64 KByte L1 cache & 32 KByte shared memory or 32 KByte L1 cache & 64 KByte shared memory. This is similar to the specifications of the Epiphany CPUs. However, the TU 102 has – compared to the Epiphany chips – a 384 bit wide memory bus with a clock rate of up to 14 gigabit per second to external GDDR6 memory Modules (672 gigabyte per second or 168 giga-words per second for 32 bit floating point numbers). It needs to be noted that these 2.3 giga-words per second of 32 bit floating point numbers per SM can only be achieved if special care is taken in correctly aligning the memory access (see CUDA Toolkit Documentation – Best Practices Guide v10.1.168 https://docs.nvidia.com/cuda/index.html).

For the timing tests in the results section, one of the used systems contains Intel Xeon E5-2640v3 CPUs. Such a CPU has 8 cores with a 2.6 GHz clock rate (in a multi-core task) and 59 gigabyte per second of memory bandwidth. A nvidia TU 102 GPU has 72 SMs. For the following calculation it is assumed that this corresponds roughly to 9 × 8 CPU cores. Following this assumption, this can be translated into 9 × Intel Xeon E5-2640v3 with a combined memory bandwidth of 531 gigabyte per second. Even though the memory on the GPU is a bit faster, the problems encountered with the multi-core CPUs can be assumed to occur with GPUs too. Even more so if the complex requirements of GPUs for memory alignment cannot be fulfilled for all 72 simulations processes at the same time.

The proposed design for the SbS IPs is optimized with respect to the number of memory accesses. The pipelined structure of the computations requires only a minimum of memory read and write transactions. For example the update of *N* latent variables in one IP requires to read 4*N* values from memory and to write *N* values back to memory. In the case of a normal CPU, basic mathematical operations are combined and for every mathematical operation transactions with the memory are required. First, for *N* sets of two numbers are multiplied which need to read from memory. Then the resulting *N* numbers need to be stored into *N* auxiliary variables within memory. The auxiliary variables are read from memory and summed up. The result of this summation is inverted and multiplied with ϵ. Again the *N* auxiliary variables are read from memory, multiplied with the former result and written back into memory. Now the *N* auxiliary variables as well as the *N* latent variables are read from memory, summed and written back into memory. These resulting *N* values are read from memory, multiplied by a constant, and written back into memory as the result of the update of the latent variables. This sums up to roughly 7*N* read operations and 4*N* write operations. In summary 5*N* memory transaction are required for the pipeline design and 11*N* memory operations for the general purpose CPU.

The question remains for which number of neurons per SbS IPs, the propose design shines. We expect that neuron numbers of 1024 and smaller are the optimal use-case. Larger number of neurons are more prone to be simulated by modern CPU with their large & densely integrated caches. Optimally, for every SbS IP with 1024 neurons we aim for a weight matrix with 100 × 1024 values. However, during the planned transfer of the presented design to an ASIC, we will evaluate if it will be beneficial to have a mixture of sizes of SbS IPs on that chip. It is expected that this question will be dominated by the area required for the supporting circuitry for the IP’s memory. It is not expected that one ASIC will be able to accommodate a whole SbS network. Thus we focus on networks of ASICs as substrate for SbS networks, where the individual ASICs communicate with spikes (i.e. the index of the firing neurons and identifier for their IP). If this approach will work out and how the details will look like in the end will be the target of our future research.

Once available the proposed ASICs will not only serve to investigate neuronal networks based on the SbS framework for AI applications where it can serve to develop both, feed foreward deep convolutional networks as well as generative models (as e.g. deterministic auto-encoders: (Ghosh et al., 2019)) but also allow investigations of large scale models of the brain (as e.g. the visual system in cortex where many retinotopic areas are mutually connected (Van Essen et al., 1992).

## Supporting information

Supplemental images and explanations

## ACKNOWLEDGMENTS

This manuscript has been released as a Pre-Print at www.biorxiv.org (Rotermund and Pawelzik, 2018).

## Notes

#### Summary of Updates

We changed the timing simulation from Matlab to C++ and added the MNIST benchmark as an example for an SbS network to the paper. We extended the discussion and tried to improve the presentation of the SbS model.

