## Supplemental images and explanations for "Massively Parallel FPGA Hardware for Spike-By-Spike Networks"

### Supplementary Material

#### 1 SUMMARIZING THE SBS ALGORITHM

The idea behind SbS networks is as follows: A network consists of several inference populations (IPs). Every one of these IPs contains a number of $N_{H,l}$ neurons (typically $N_{H,l} \in [2, ..., 1000]$ and $l$ identifies the IP) and with every neuron $i$ there comes a latent variable $h_l(i)$. The latent variable describes the state of the neuron and can have only values in the range of $0 \leq h_l(i) \leq 1$. The neurons within an IP are in a competition. The competition is mediated via a common normalization ($\sum_i h_l(i) = 1$) over all the $h_l(i)$ belonging to that IP. Beside the normalization, there is no information exchanged within the IP. However, information is exchanged among populations via spikes $s_O^t$, where $t$ represents the time step in which the spike was emitted and $O$ referring to its origin. The origin of that spike could be from an input population which represent an input pattern $p_g(s)$ (e.g. pixel images, waveforms, time series) presented to the network. We denote the probability distribution used by the input population $g$ with $p_g(s)$, where $s$ enumerates the $N_S$ neurons in that input population. Another origin for spikes can be found in other IPs. In this case their latent variables $h_l(i)$ are interpreted as probability distributions which are used to determine which neuron will spike next. Algorithmically, generating a spike translates into drawing a random number $R \in [0, \dots, 1]$ , comparing $R$ to the cumulative sum $C(i) = \sum_{j=1}^{i} h_l(j)$ if the spike will be produced by an IP or $C(i) = \sum_{j=1}^{i} p_g(j)$ if the spike is produced by an input population. Then the $i$ for which the cumulative sum $C(i)$ surpasses $R$ is selected as spike $s_O^t$.

The emitted spikes $s_O^t$ are sent to connected IPs $l$. Which IP $l$ is listening to which spike emitting population $O$, is predetermined by the architecture of the network. If there is a connection between two populations, then for every neuron $i$ in the receiving IP $l$ there is a weights $p^{O \to l}(s_O|i)$. $s_O$ denotes the neurons in the emitting population $O$. Weight values are only from the range $p^{O \to l}(s_O|i) \in [0, 1]$. Furthermore, the weights are normalized according to $\sum_{s_O} p^{O \to l}(s_O|i) = 1$.

In (Ernst et al., 2007; Rotermund and Pawelzik, 2019a,b) we presented the updating rules for the latent variables and learning rules for the weights in much detail, including their derivation. In the following the idea and origin behind the algorithms are only briefly recapitulated: The core of the IP is a generative model using non-negative entities. The latent variables of the IP $h(i)$ are used in accordance with the weights $p(s|i)$ to represent a reconstruction of the input $p_\mu(s)$ ($\mu$ enumerated the input pattern from a ensemble of $N_M$ patterns) to the IP via:

$$r_\mu(s) = \sum_{j}^{N_H} p(s|j) h_\mu(j) \tag{S1}$$

The goal is to approximate $p_\mu(s)$ with $r_\mu(s)$ as well as possible. The distance is measured with the cross-entropy

$$E = -\sum_{\mu}^{N_M} \sum_{s}^{N_S} p_\mu(s) \log\left(r_\mu(s)\right) \tag{S2}$$

Derivatives of the cross-entropy are calculated for getting the gradients for $p(s'|j')$ and $h_{\mu'}(i')$:

$$-\frac{\partial E}{\partial h_{\mu'}(i')} = \sum_{s}^{N_S} p_\mu(s) \frac{p(s|i')}{\sum_j^{N_H} p(s|j) h_{\mu'}(j)} \tag{S3}$$

$$-\frac{\partial E}{\partial p(s'|j')} = \sum_{\mu}^{N_M} p_\mu(s') \frac{h_\mu(i')}{\sum_j^{N_H} p(s'|j) h_\mu(j)} \tag{S4}$$

Concerning the update rule of the latent variables, we implement three changes: 1.) Input distribution $p_\mu(s)$ is replaced by only the next spike $s^t$ ($p_\mu(s) = \delta_{s,s^t}$). 2.) We make a multiplicative gradient update out of $\frac{\partial E}{\partial h_{\mu'}(i')}$ by multiplying it with $\epsilon(t) h_\mu(i')$, where $\epsilon(t)$ is an update parameter which can be modulated over the number of processed spikes. $\epsilon$ acts like a low pass filter which smoothes the latent variables when they are updated with the next spike. 3.) The updated latent variables need to be normalized. Thus the normalization is ensured by a multiplicative factor $\frac{1}{\epsilon(t)}$. All three steps, result in

$$h_\mu^{t+1}(i) = \frac{1}{\epsilon(t)} \left( h_\mu^t(i) + \epsilon(t) \frac{p(s^t|i) h_\mu^t(i)}{\sum_j^{N_H} p(s^t|j) h_\mu^t(j)} \right) \tag{S5}$$

as update rule for the latent variables of an IP with incoming spike $s^t$.

Based on the gradient $\frac{\partial E}{\partial p(s'|j')}$, it is possible to derive four local learning rules. A multiplicative learning rule

$$\tilde{p}^{l+1}(s|j) = p^l(s|j) + \gamma(l) \sum_{\mu}^{N_M} p_\mu(s) \frac{p^l(s|i) h_\mu(i)}{\sum_j^{N_H} p^l(s|j) h_\mu(j)} \tag{S6}$$

and an additive learning rule

$$\tilde{p}^{l+1}(s|j) = p^l(s|j) + \gamma(l) \sum_{\mu}^{N_M} p_\mu(s) \frac{h_\mu(i)}{\sum_j^{N_H} p^l(s|j) h_\mu(j)} \tag{S7}$$

using (mini-) batches of input patterns. $\gamma(l)$ is a learning rate that can change with the number of already realized weight updates. Furthermore, online learning rules are possible which update the weights with each incoming spike $s^t$. Here also a multiplicative learning rule

$$\tilde{p}^{l+1}(s|j) = p^l(s|j) + \gamma(l) \delta_{s,s^t} \frac{p^l(s|i) h_\mu(i)}{\sum_j^{N_H} p^l(s|j) h_\mu(j)} \tag{S8}$$

and an additive version

$$\tilde{p}^{l+1}(s|j) = p^l(s|j) + \gamma(l) \delta_{s,s^t} \frac{h_\mu(i)}{\sum_j^{N_H} p^l(s|j) h_\mu(j)} \tag{S9}$$

are available. All these learning rules don't ensure the normalization of the weights after the learning step. Thus re-normalization via

$$p^{l+1}(s|j) = \frac{\tilde{p}^{l+1}(s|j)}{\sum_{s'}^{N_S} \tilde{p}^{l+1}(s'|j)} \quad \text{(S10)}$$

is required. In Rotermund and Pawelzik (2019a) we also presented another batch learning rule that is non-local in nature and based on the idea of error back-propagation, somewhat similar to learning rules used in many non-spiking neuronal networks. In the case of an input population $p_\mu(s)$ represents an input probability distribution which is given by the input pattern. If the spikes are received from another IP $g$, then $p_\mu(s)$ is given by the latent variables of that IP $h_g^t(s)$, which were also used for generating the observed spike $s^t$.

#### 2 ADDITIONAL FIGURES AND TABLES

##### 2.1 Non-negative numbers

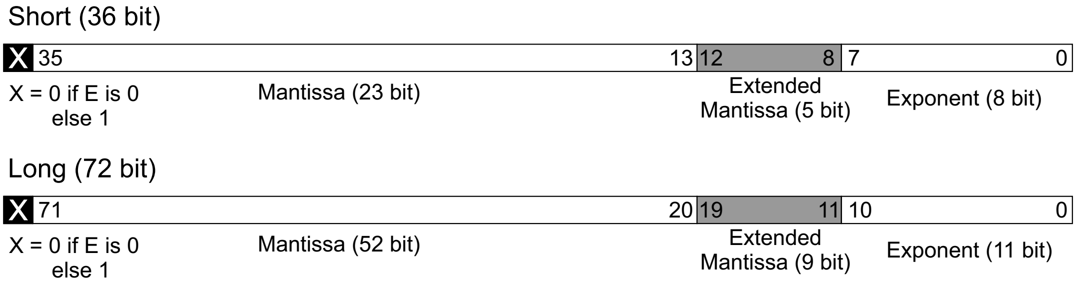


**Figure S1.** Representation of custom non-negative floating point number with 36 bits and 72 bits. The most significant bit of the mantissa is shown as $X$ in the black box and is not stored in memory. $X$ is set to 0 if the exponent is zero and set to 1 otherwise. The bits shown in gray are the extension to the precision for the mantissa over the IEEE 747 standard for single or double precision.

#### 2.2 Computational building blocks

| | | Floating point | Fixed point |
|---|---|---|---|
| **Module** $*$ | LUT | 1425 | 442 |
| | LUT-FF pairs | 1674 | 614 |
| | Slice registers | 1844 | 676 |
| **Module** $+$ | LUT | 433 | 20 |
| | LUT-FF pairs | 358 | 20 |
| | Slice registers | 429 | 0 |
| **Module** $\#$ | LUT | 2407 | 1333 |
| | LUT-FF pairs | 2594 | 1409 |
| | Slice registers | 2992 | 1750 |
| Operation $1/X$ seq. | LUT | 301 | 197 |
| | LUT-FF pairs | 304 | 311 |
| | Slice registers | 213 | 180 |
| **Module MEM** | LUT | 383 | 345 |
| | LUT-FF pairs | 728 | 562 |
| | Slice registers | 816 | 661 |
| | BRAM $h(i)$ | 1x 36k | 1x 18k |
| | BRAM $p(s\|i)$ | 1x 36k | 1x 18k |
| **Module NORM-MULTI** | LUT | 3306 | 1345 |
| | LUT-FF pairs | 3916 | 1501 |
| | Slice registers | 4349 | 1503 |
| Operation $\sum$ | LUT | 452 | 48 |
| | LUT-FF pairs | 496 | 60 |
| | Slice registers | 569 | 41 |
| Operation $Y/X$ seq. | LUT | 426 | 322 |
| | LUT-FF pairs | 457 | 324 |
| | Slice registers | 340 | 227 |

**Table S1.** Basic computational building blocks: Components used for the building block. All addresses are represented by 10 bit integer values. The BRAM counts for **module MEM** are given for up to 1024 $h(i)$ values and up to 1024 $p(s|i)$ values. For the fixed point modules, 18 bit integers are used as input. The operations are sub-parts of modules and are listed as extra detailed information.

| | | Floating point | Fixed point |
|---|---|---|---|
| **Module** $*$ | clock cycles | 13 | 8 (42bit) |
| **Module** $+$ | clock cycles | 8 (36bit), 13 (72bit) | 1 |
| **Module** $\#$ | | | |
| (calculate $\frac{1}{1+A}$) | clock cycles | 47 | 46 (23bit) |
| (multiply with $B$) | clock cycles | 14 | 10 (23bit) |
| Operation $1/X$ seq. | clock cycles | 34 | 39 (24bit) |
| **Module MEM** (and $W(S\|i)$) | clock cycles | 14 | 14 |
| **Module NORM-MULTI** | | | |
| ($\epsilon$/norm, after last value) | clock cycles | 97 | 61 ($\epsilon$ 22bit) |
| (multiplication pipeline) | clock cycles | 15 | 11 |
| Operation $\sum$ (after last value) | clock cycles | 39 | 4 (22bit) |
| Operation $Y/X$ seq. | clock cycles | 35 | 45 (22bit) |

**Table S2.** Basic computational building blocks: Number of clock cycles between the input into the module and when the output of the computation is available. For the **module** $\sum$ and where it is used in **module NORM-MULTI**, the number of clock cycles after when the last input values was presented are shown. Since the numbers are measured in circuit, deviations from the 18bit fixed point input into the modules are noted in brackets.

| | | Floating point | Fixed point |
|---|---|---|---|
| **Module Spike Generator** (fig.S2a) | LUT | 234 | 82 |
| | LUT-FF pairs | 274 | 162 |
| | Slice registers | 257 | 116 |
| **Module Spike Generator with offset** (fig. S4a) | LUT | 647 | 231 |
| | LUT-FF pairs | 1137 | 614 |
| | Slice registers | 1158 | 569 |
| (after threshold) | clock cycles | 13 | 9 |
| Mersenne twister (MT) 19937 (fig. S3) | LUT | 1326 | 1326 |
| (produces 32 bit random number) | LUT-FF pairs | 1474 | 1474 |
| | Slice registers | 1137 | 1137 |
| | BRAM | 2x 36k | 2x 36k |
| **Module Rate Calculator** (fig.S2b) | LUT | 2776 | 854 |
| | LUT-FF pairs | 3114 | 1000 |
| | Slice registers | 3527 | 957 |
| | BRAM | 1x 36k | 1x 36k |
| (calculate $\frac{1}{C_{All}}$) | clock cycles | 51 | 48 |
| (recall $\hat{p}(s)$) | clock cycles | 26 | 14 |
| **Module Normalized Weight Offset** (fig. S4b) | LUT | 3289 | 1859 |
| | LUT-FF pairs | 3771 | 2067 |
| | Slice registers | 4165 | 2358 |
| (calculate $\frac{1}{1+\varphi}$, $\frac{\varphi}{N_S}$) | clock cycles | 49 | 58 |
| (operate) | clock cycles | 25 | 15 |

**Table S3.** Number of components as well as number of clock cycles for the modules concerned with generating spikes, producing random numbers (via Mersenne twister 19937), calculating rates out of observed spikes, and adding a normalized offset to the weights during learning. The number of clock cycles listed for the spike generation module denotes how long it takes after the input value responsible for producing the spike in the end (i.e. exceeding the threshold by the cumulative sum) is observed.

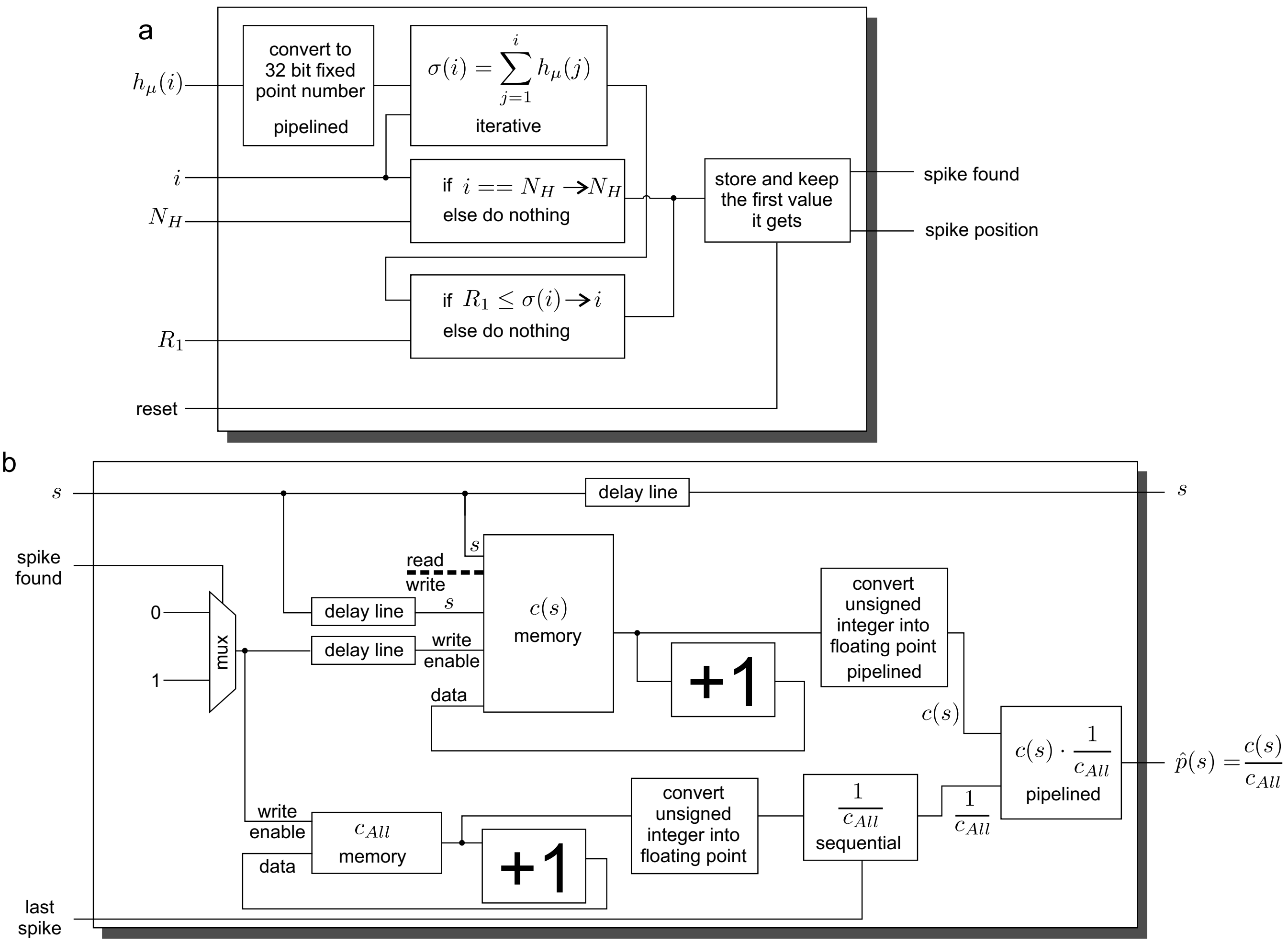


**Figure S2.** a.) Module for generating spike from a random number and probability distribution. b.) Module for calculating a rate from observed spikes. The conversion from unsigned integer into floating point number is only used in the floating point version.

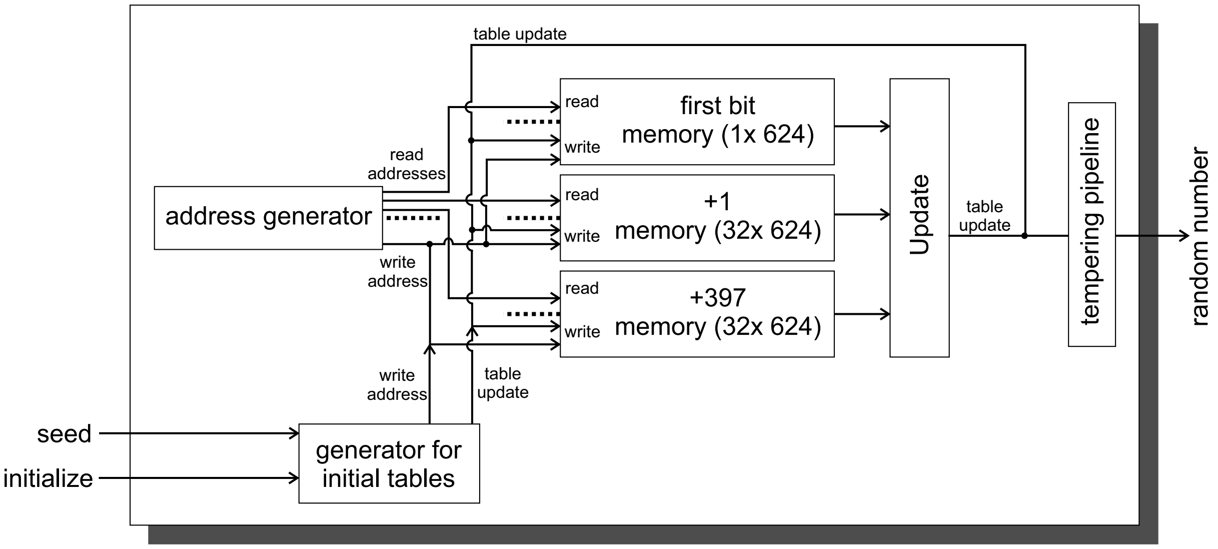


**Figure S3.** Overview for the Mersenne twister (MT 19937) module.

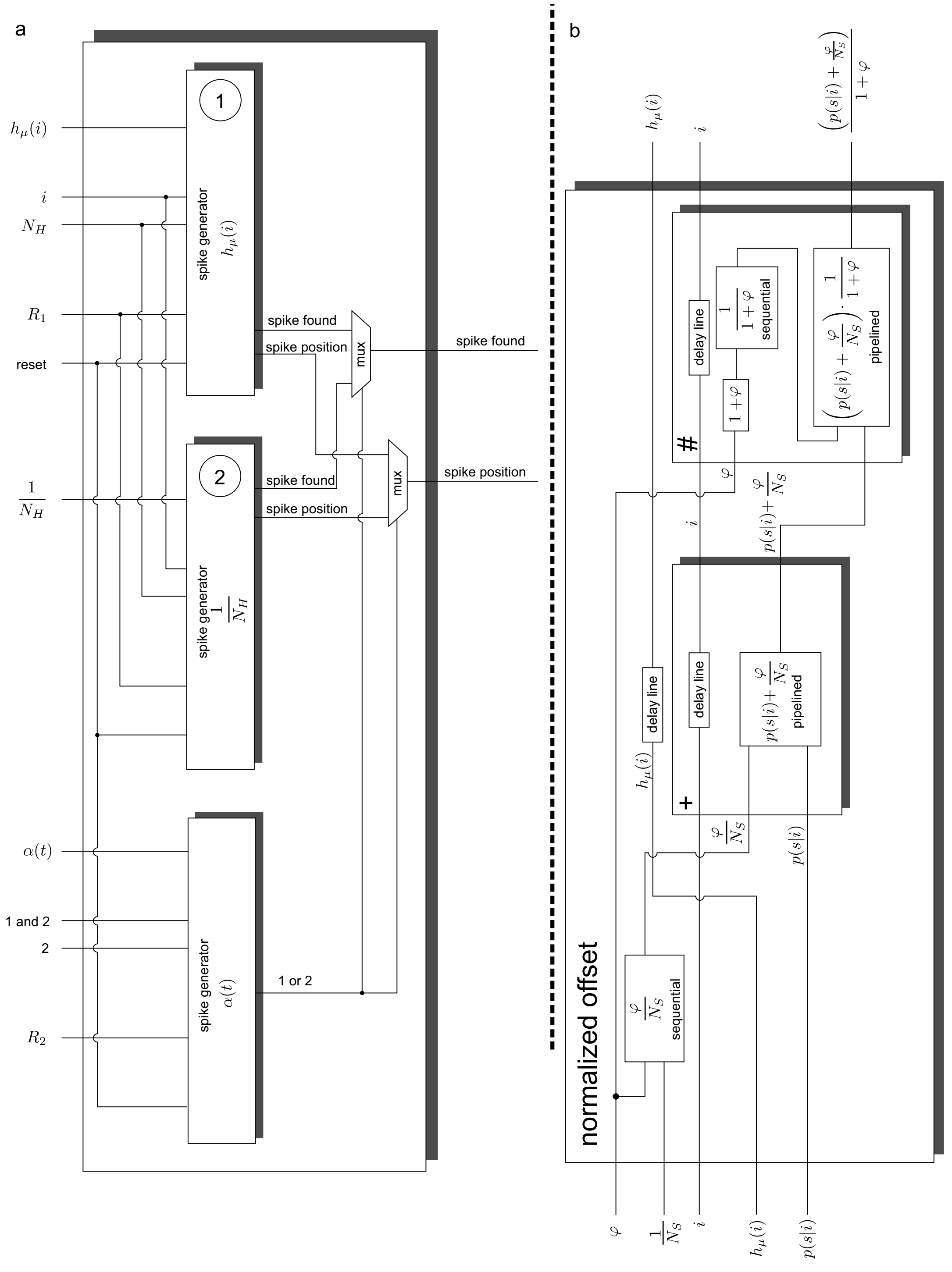


**Figure S4.** a.) Module for generating spikes with an uniform offset. b.) Module for adding a normalized offset to weights.

#### 2.3 Circuits for updating $h(i)$, online and batch learning $p(s|i)$

| | | Floating point | Fixed point |
|---|---|---|---|
| h dyn. plus p online lear. (2. stage) | LUT | 6437 | 5296 |
| | LUT-FF pairs | 6908 | 5670 |
| | Slice registers | 8285 | 7020 |
| (h update, latency) | clock cycles | 39 | 23 |
| (p update, latency) | clock cycles | 15 | 10 |
| p online learning (3. stage) | LUT | 2857 | 1520 |
| | LUT-FF pairs | 3293 | 1787 |
| | Slice registers | 3831 | 2077 |
| **Module Memory** $U(i)$ | LUT | 33 | 9 |
| | LUT-FF pairs | 147 | 155 |
| | Slice registers | 143 | 162 |
| | BRAM | 1x 36k | 1x 36k |
| (recall latency) | clock cycles | 5 | 5 |
| p batch learning (2. stage) incl. $W$ | LUT | 3329 | 1374 |
| | LUT-FF pairs | 3848 | 1619 |
| | Slice registers | 5000 | 2100 |
| | BRAM | 2x 36k | 1x 36k |
| **Module Memory** $W(s|i)$ | LUT | 459 | 370 |
| | LUT-FF pairs | 644 | 508 |
| | Slice registers | 767 | 612 |
| | BRAM | 2x 36k | 1x 36k |

**Table S4.** Number of components and clock cycles for the circuits for updating $h(i)$, online and batch learning $p(s|i)$.

#### 2.4 Connecting Spike-By-Spike inference populations and input populations

| | | Floating point | Fixed point |
|---|---|---|---|
| SbS inference population with 2 connection lists | LUT | 26915 | 16794 |
| | LUT-FF pairs | 32508 | 20822 |
| | Slice registers | 40239 | 25317 |
| | BRAM | 12x 36k | 9x 36k, 2x 18k |
| SbS arithmetic core | LUT | 23779 | 13914 |
| | LUT-FF pairs | 27534 | 16004 |
| | Slice registers | 33965 | 19443 |
| | BRAM | 6x 36k | 3x 36k, 2x 18k |
| Connection list (32 entries) | LUT | 1475 | same |
| | LUT-FF pairs | 2019 | same |
| | Slice registers | 2829 | same |
| | BRAM | 3x 36k | same |
| **Broadcast module** | LUT | 25 | same |
| | LUT-FF pairs | 33 | same |
| | Slice registers | 29 | same |
| Input population | LUT | 407 | same |
| | LUT-FF pairs | 548 | same |
| | Slice registers | 605 | same |
| | BRAM | 1x 36k | same |

**Table S5.** Component counts for the modules shown in figure 6.

#### 2.5 Coordinating the networks and exchanging data

| Type | | |
|---|---|---|
| **Message control center** | LUT | 2164 |
| | LUT-FF pairs | 2703 |
| | Slice registers | 2554 |
| | BRAM | 8x 36k |
| MT with message interface | LUT | 1536 |
| | LUT-FF pairs | 1650 |
| | Slice registers | 1475 |
| | BRAM | 2x 36k |
| Arbiter (databus incoming) | LUT | 162 |
| | LUT-FF pairs | 170 |
| | Slice registers | 119 |
| Arbiter (databus outgoing) | LUT | 311 |
| | LUT-FF pairs | 459 |
| | Slice registers | 424 |

**Table S6.** Component count for the modules on the 32 bit data bus.

#### 2.6 Pipeline designs for the floating point operations

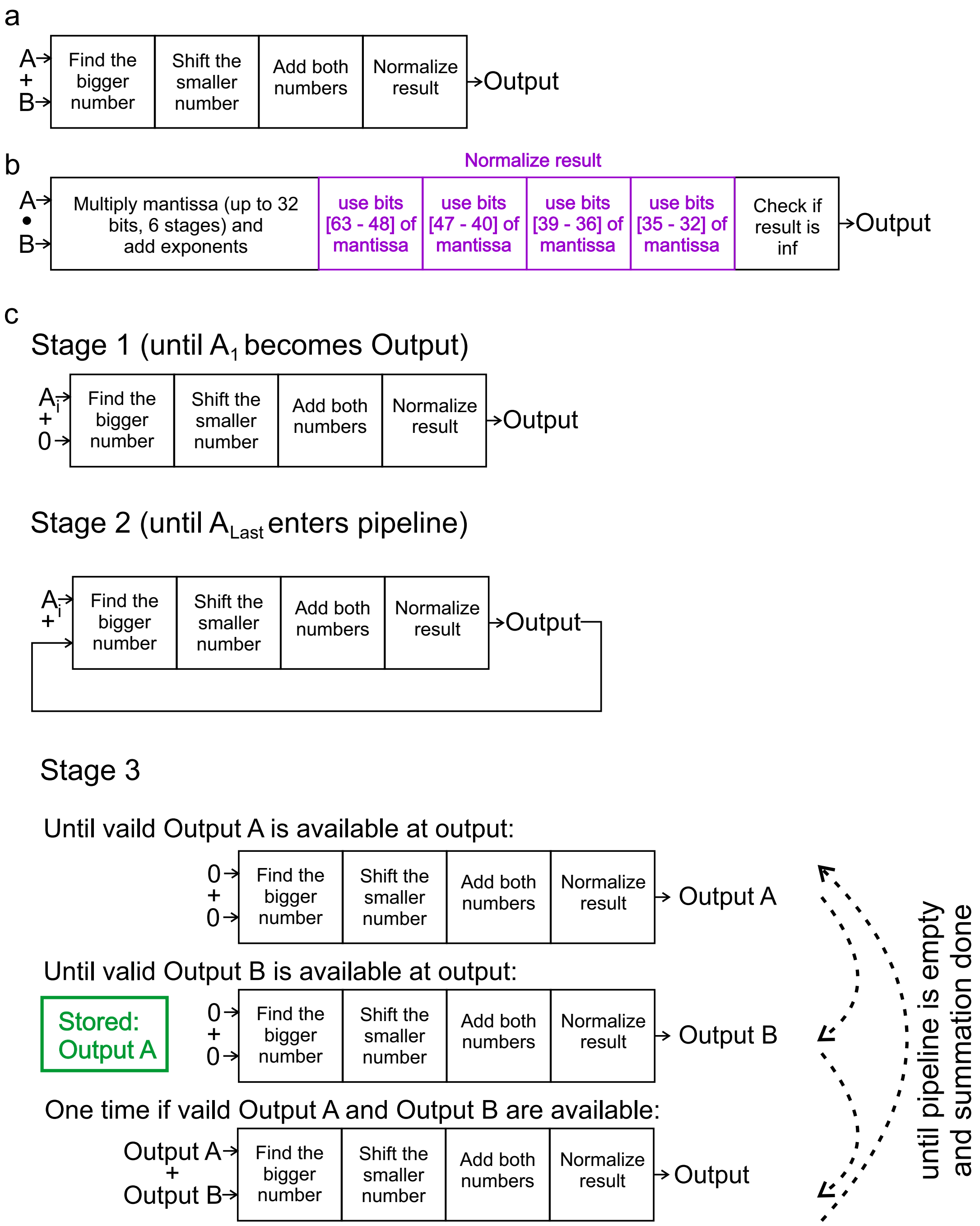


**Figure S5.** Pipeline designs used for a) adding two non-negative 36 bit floating point numbers and b) multiplying two non-negative 32 bit numbers. The buffering steps of auxiliary results for improving the timing are not shows. c) Three stage pipeline design for calculating a cumulative sum from a given series of $A_i$'s.

#### 2.7 Comparing CPU with FPGA

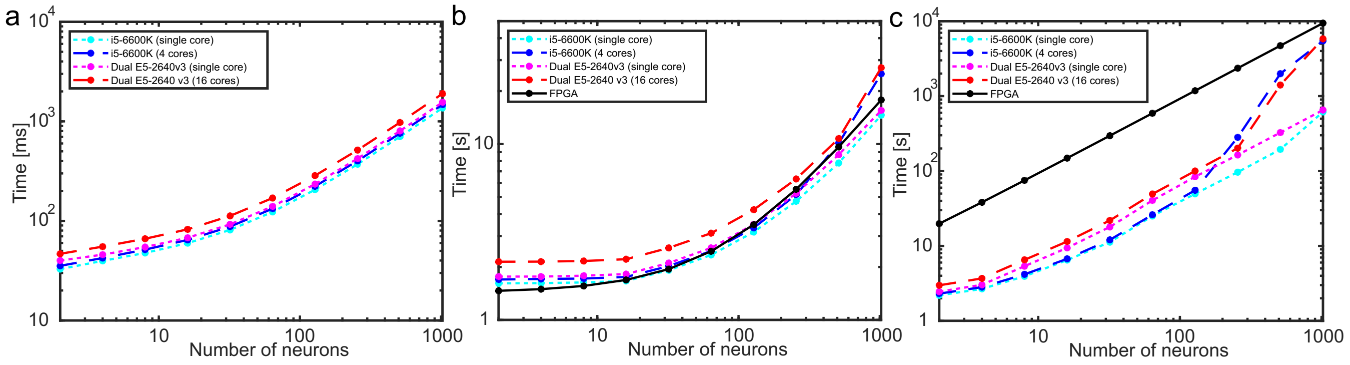


**Figure S6.** a.) Time for drawing one million spikes from a random probability distribution with the number of neurons shown on the x-axis. If $N_H$ is large enough, the FPGA does not depend on $N_S$, the generation of spikes is done by in parallel to other operations, and doesn't require additional clock cycles for this operation. b.) Duration for updating the latent variables with one million spikes. The input population was kept at $N_S = 1024$. $N_H$, the number of neurons in one SbS inference population, was varied according to the x-axis. The Intel Core i5-6600K was running as single core and used Turbo Boost mode with 3.8GHz. c.) Durations for the simulation of b.) with active online learning and adding a normalized offset to the weights. The shown durations are averaged over 100 trials times the number of used cores. The black lines represents the estimated time which the FPGA needs.

In the following, we will compare the performance of general purpose CPUs (Intel Core i5-6600k with four cores and dual Intel Xeon E5-2640v3 with eight cores each) with the hardware implementation of the SbS IPs. Details concerning the applied methods and the used source code are in the supplemental materials.

In figure S6a, the time for drawing one million spikes from a random probability distribution on a single CPU core is shown. In comparison, the FPGA requires roughly 10 clock cycles (80ms for one million spikes) plus the number of probability values $p(s)$ in clock cycles until the spike was found. As long as $N_S + 10 - (179 + 2 \cdot N_H)$ is smaller or equal zero, no additional time is required and the spike is produced in parallel to updating the latent variable. Otherwise only $N_S + 10 - (179 + 2 \cdot N_H)$ additional clock cycles are required. For the CPUs, the time required for producing these spikes scales also linearly with $N_S$. Without a parallel update of the latent variable which absorbs clock cycles, the FPGA would be ≈4.2x slower ($N_S = 1024$) in drawing spikes that the Intel Core i5 in Turbo Boost Mode.

Figure S6b shows the duration it takes to update the latent variables one million times on the Intel Core i5 and the FPGA. All durations reflect the time, one SbS inference population needs to finish the task. For SbS inference populations with small numbers of neurons (below 128 neurons), the FPGA is slightly faster. For larger number of neurons the CPUs are faster if they are used in single core mode but slower when multi cores are used. In the case of $N_H = 1024$, the FPGA requires 17.8s while the i5 (14.6s) and the E5 (15.5s) when used in single core mode are a bit faster. If the CPUs are operated in multi core mode, the i5 needs 25.1 s and the E5 requires 27.2s, which is a bit slower than the FPGA. If the intended speed of 250MHz for the FPGA could be realized, the duration the FPGA requires would be halved. Furthermore, the FPGA also performs an additional but optional normalized offset to the weights in the reported duration, which we didn't include in these C++11 simulations.

Concerning online learning, the FPGA can calculate part of the learning rule in parallel to the update of the latent variable. However, the re-normalization of the updates weights takes a substantial amount of time ($N_H \cdot (124 + N_S)$ clock cycles or 1175552 clock cycles / 9.4ms for one re-normalization with $N_H = 1024$ and $N_S = 1024$). Due to the high clock rates and the simple calculations, Intel CPUs are able to perform this calculation very fast as long as the memory bandwidth is not hit. Figure S6c shows that for single core operation with intermediate number of neurons per SbS inference population, the Intel CPUs are roughly 20-23x faster than the FPGA. For larger number of neurons, the memory bandwidth starts to get a bottleneck, where a reduction to ≈14x can be seen. For smaller number of neurons, the performance is only 8x faster than the FPGA because the CPU can't simply stream the data from the memory to it's floating point units but is interrupted by other operations. Using more than a single core shows the full size of the problem with the memory bandwidth limit where the performance for larger number of neurons breaks down to a speed increase of only ≈1.6x over the FPGA. This effect doesn't occur on the FPGA since every SbS inference population has it's independent memory. Nevertheless, in the case with $N_H = 1024$, 26 SbS inference populations on a FPGA with a 125MHz clock would be required to reach performance equality with the 16 core dual Intel Xeon system.

In the case of batch learning, the required time is almost exclusively defined by the update of the latent variables. The calculation of $W(s|i)$ from the $h(i)$, $p(s|i)$ and $\hat{p}(s)$ is performed only once per pattern after many spikes have been processed.

#### 2.8 MNIST example with fixed point numbers

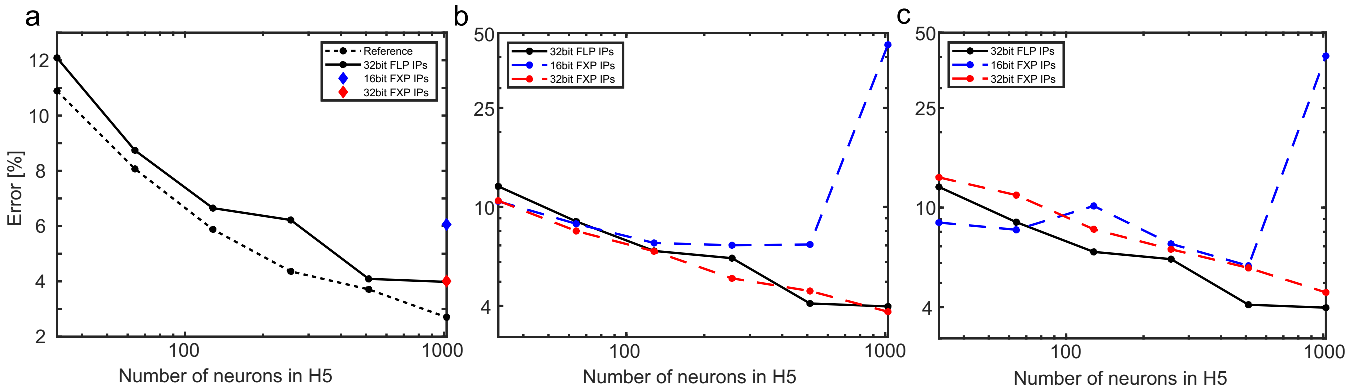


**Figure S7.** Classification errors for the tested variants of the MNIST SbS network in dependency of the number of neurons (32, 64, 128, 512, and 1024) in the $H5$ IP. a.) As reference (black dotted line), the performance for a network, which uses everywhere 32 bit floating point (32 bit FLP) IPs, is shown. The black solid line, which reappears in subplot b.) and c.), uses 16 bit fixed point (16bit FXP) IPs for the layers $H1$ to $H4$ while the layers $H5$ and $HY$ are 32 bit FLPs. The blue and red diamonds show the error when the weights from the 32 FLP IPs case (black solid line) with 1024 $H5$ neurons are converted into 16 bit / 32 bit FXP weight sets and used with their respective FXP IPs. For the results shown in b.) and c.) the layers $H1$ and $H4$ use also the 16bit FXP IPs. b.) The dynamics of the latent variable in layers $H5$ and $HY$ are simulated using 16bit FXP IPs for the blue curve and 32bit FXP IPs for the red curve. For learning, first the values stored in the latent variables are converted into 32bit FLP, then the weights are updated, and finally converted into 16 bit / 32 bit FXP weights for using them in the dynamic of the latent variables. c.) Like b.) but learning is performed fully with FXP weights and calculations.

For testing the influence of using fixed point number in SbS networks, we examined different versions of representation for these non-negative numbers through Matlab simulations, using custom mex C extensions. For measuring the differences in performance, we used a simplified version of the MNIST network from (Rotermund and Pawelzik, 2019a) (shown in figure 1). This network was trained to classify the digits shown in images (28 x 28 pixel with 8 bit gray values for the pixels) of handwritten digits. The MNIST benchmark data base contains 60,000 training pattern and 10,000 test patterns. We used the shown SbS network only in a feed-forward fashion. The network, the applied learning procedure, and more information about the simulation can be found in the supplemental materials.

The network consists of: 1.) An input layer $X$, with 24 x 24 input populations with 50 neurons each. 2.) The first convolution layer $H1$ which uses 5 x 5 spatial kernels (with stride 1 and no zero padding) and contains 24 x 24 SbS inference populations (IPs) with 32 neurons each. 3.) The first pooling layer $H2$, which combines 2 x 2 spatial activity patches (with stride 2) from $H2$ into one IP each. $H2$ has 12 x 12 IPs with 32 neurons each. 4.) The second convolution layer $H3$, which like $H1$, has 5 x 5 spatial kernels (with stride 1 and no zero padding) but with 64 neurons each. 5.) The second pooling layer $H4$, which combines 2 x 2 spatial activity patches (with stride 2) from $H3$ into one IP each. $H4$ has 4 x 4 IPs with 64 neurons each. 6.) A fully connected layer $H5$ with 1 IP which has e.g. 1024 neurons (this number of neurons will be changed for different simulations). 7.) The output layer $HY$ which is one IP with 10 neurons. Every one of these 10 neurons is associated to one of the ten classes of digits, during learning. For decoding the result of the computation, the output neuron is selected with the highest value in their latent variables and its associated digit class is used.

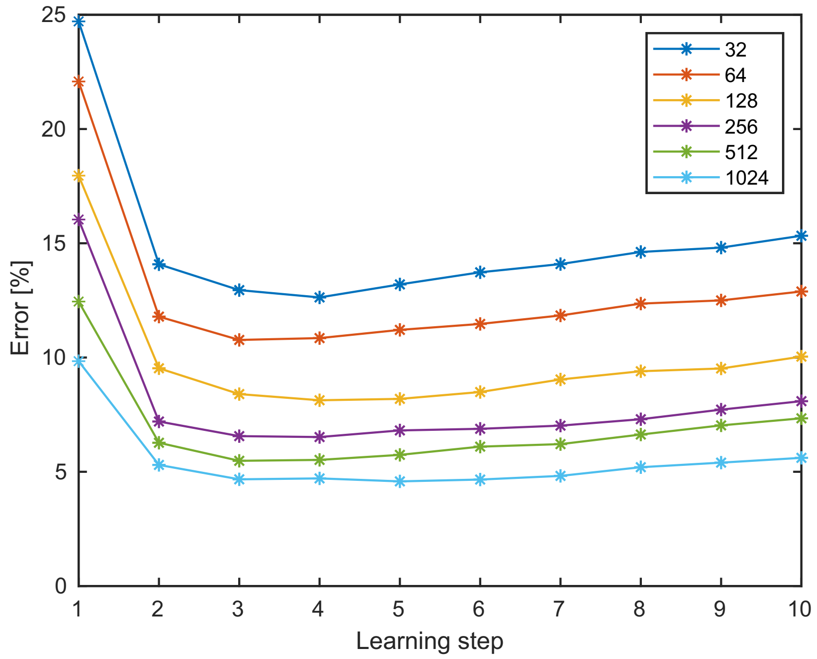


**Figure S8.** Development of the error in dependency of the number of the learning steps for the 32 bit FXP case shown in figure S7c.

(Rotermund and Pawelzik, 2019a) details how this network is set up and trained. In the following only the stylized facts are presented: a.) Every input population and IP produces one spike each per time step of the simulation. b.) The IPs of the pooling layers use the same update rule for the latent variables as all other IPs too. For turning them into pooling layers pre-defined fixed weight matrices are used. These weights matrices ensure that the spikes produced by the 32 neurons (each of these neurons represents one so called 'feature' of the IPs input) from $H1$ IPs (or by the 64 neurons from $H3$ IPs) are processed such that features are spatially combined but are in a competition with different features. This ensures that the strongest features survive while the weaker ones are suppressed. c.) The weights matrices of the convolution layers are shared for all 24 x 24 spatial positions in $H1$ (and 8 x 8 positions in $H3$). d.) For keeping the computational requirements low, the 28 x 28 pixel input images are converted into 24 x 24 patches with 5 x 5 pixels. This emulates the convolution process of $H1$ by performing the convolution on the input side rather in convolution layer. The 5 x 5 pixel patches are further pre-processed by performing an On/Off cells transformation. This corresponds to that the images' 8 bit unsigned integer values are transformed into values from the $\pm 1.0$ range by removing the mean and re-scaling the pixel values to the new value range. Then the positive and the negative numbers are separated into two different non-negative channels (hence half of the channels will contain the value 0 by construction). Thus the 5 x 5 pixel patches are converted into 5 x 5 x 2 patches containing non-negative numbers. By applying the L1 norm, this results in 24 x 24 input probability distributions with 50 values each which are used for producing the spikes for the input layer.) e.) During simulating the network for one given input image, 1200 spikes are produced by every individual IP and input population. For the first 1000 spikes $\epsilon(t) \approx 0.1$ and for the latter 200 spikes $\epsilon(t) \approx \frac{0.1}{25}$. The reduction in $\epsilon$ helps to smooth fluctuations out of the latent variables. The $\approx$ stems from converting floating points into unsigned integers for the fixed point types of SbS IPs.

For learning the equations for multiplicative batch learning was used:

$$\tilde{p}^{l+1}(s|j) = \gamma_1 p^l(s|j) + \gamma_2 \sum_{\mu}^{N_M} p_\mu(s) \frac{p^l(s|i) h_\mu(i)}{\sum_j^{N_H} p^l(s|j) h_\mu(j)} \tag{S11}$$

$$p^{l+1}(s|j) = \frac{\tilde{p}^{l+1}(s|j)}{\sum_{s'}^{N_S} \tilde{p}^{l+1}(s'|j)} \tag{S12}$$

The networks are trained in a layer by layer fashion (see pre-learning in (Rotermund and Pawelzik, 2019b) for more details). First the shared weights between the $H1$ are trained by 20 learning steps with presenting all training data for ever learning step. These weights are used in an additional simulation to propagate the input pattern into $H1$ latent variable distributions. Then the pooling layer $H2$ is used in another simulation to produce $H2$ latent variable distributions. After that the weights between pooling layer $H2$ and convolution layer $H3$ are trained for 20 learning steps. These weights are used to propagate the $H2$ latent variable distributions into $H3$ activity and then – via pooling layer $H4$ – into $H4$ latent variable distributions. This was done for two types of networks: a.) With IPs that used 16 bit fixed point numbers (16 bit FXP) for latent variables $h(i)$ and weights $p(s|i)$ ($\gamma_1 = 0$ and $\gamma_1 = 1.0$). In this case, we represent non-negative real numbers for the value range $[0, ..., 1]$ via 16 bit integers through $X = \frac{X_{Int}}{2^{16}-1}$. For preventing loss of accuracy, we extended the number of bits for intermediate results, like it was also done for the hardware design. b.) The weights and latent variables are represented by 32 bit floating point number (32 bit FLP) in the Matlab simulations ($\gamma_1 = 0$ and $\gamma_2 = 1.0$). We used 64 bit floating point numbers (IEEE 754, double precision) in the Matlab simulations for collecting all the contributions from the 60,000 training patterns. c.) During training, after every learning step a small number is added to keep the weights values away from a perfect zeros. We used $\frac{0.05}{N_S}$ for everything learned with 32 floating point numbers. When learning with 16 bit fixed point numbers, the value 3 was used for the weights between $H4$ and $H5$ as well as 33 for the weights between $HY$ and $H5$. For the 32 bit fixed point case, 768 and 8448 were used respectively.

For the weights between $H4$ and $H5$ as well as $H5$ and $HY$, the learning procedure was modified. ($HY$ is defined during training by the associated digit class for the input used for layer $X$.) During training, layer $H4$ and layer $HY$ are combined and used as input for layer $H5$. Then the weights between the combined layer $H4\&HY$ and $H5$ are learned for a given number of learning steps. After that the resulting weights are separated which results in the required weights between layer $H4$ and $H5$ as well as layer $H4$ and layer $HY$. Except the 32 bit FXP case in figure S7c in the mai npaper where 5 learning steps for these weights were used, all curves in figure S7 in the main paper are based on weights after 20 learning steps.

In more detail: $I_{\mu,x,y}(s)$ represents the input distribution which is used to generate the spikes that are sent to the normalization group in $H1$ with their $h_{\mu,x,y}(i)$ (from now on denoted by $h_{H1,\mu,x,y}(i)$ to keep track of the layers). $\hat{I}_{\mu,x,y}(s)$ is calculated by counting the input spikes drawn from $I_{\mu,x,y}(s)$ and then converting it into a probability distribution by normalization. Beside of averaging over all 60000 patterns $\mu$, the contributions from all the 24 x 24 spatial positions are averaged too. Learning is repeated for 20 steps. After that the network is simulated a 21th time and the resulting $h_{H1,\mu,x,y}(i)$ are stored. Now this collection of 60000 $h_{H1,\mu,x,y}(i)$ is used as input $I_{H1,\mu,x,y}(s)$ for a second one layer network with layer $H2$ ($h_{H2,\mu,x,y}(i)$) as hidden layer. Running this network once with the fixed pooling weights propagates the input $I_{H1,\mu,x,y}(s)$ one layer further to $h_{H2,\mu,x,y}(i)$. Again we use $h_{H2,\mu,x,y}(i)$ as $I_{H2,\mu,x,y}(s)$ in a third one layer network where the hidden layer is $H3$. These weights are trained for 20 learning steps and then with the 21th run of the simulation, the input $I_{H3,\mu,x,y}(s)$ is calculated. $I_{H3,\mu,x,y}(s)$ is propagated further via pooling layer $H4$ with its fixed weights into $I_{H4,\mu,x,y}(s)$. Up to this layer the learning procedure was unsupervised.

For the last two layers $H5$ and $HY$, learning is slightly modified. $I_{H4,\mu,x,y}(s)$ defines the input into $H4$ and the digit for a given pattern defines $h_{HY,\mu,x,y}(i)$. All $h_{HY,\mu,x,y}(i)$ are zero except the one neuron which is associated to the digit. This neuron's $h_{HY,\mu,x,y}(i)$ is set to one. For training, like shown in Ernst et al. (2007), such a two layer network can be converted into a one layer network by folding down $h_{HY,\mu,x,y}(i)$ and using it as additional input $I_{HY,\mu,x,y}(s)$ to $H5$. In every time step, we now draw 4 x 4 spikes from $I_{H4,\mu,x,y}(s)$ and one spike from $I_{HY,\mu,x,y}(s)$. This allows us to train the weights $W^{H4\rightarrow H5}$ and $W^{HY\rightarrow H5}$ simultaneously for 20 learning steps like it was done for the other layers before. However, for testing the performance of this network the weights $W^{H5\rightarrow HY}$ are required. We calculate them from $W^{HY\rightarrow H5}$ by transposing the matrix and re-normalizing it afterwards.

Figure S7a, shows as reference the error in a network which consists only of 32 bit FLP IPs ($\gamma_1 = 0$ and $\gamma_2 = 1.0$). In comparison, the error of a network where layer $H1$ to $H4$ use 16 bit FXP and only layer $H5$ and $HY$ use 32 bit FLP IPs ($\gamma_1 = 0$ and $\gamma_2 = 1.0$) is shown as solid black line in figure S7a. The performance is moderately reduced. Using the corresponding weights for 1024 neurons (last point of the solid black line), converting these FLP weights into 16 bit / 32 bit FXP weights, and running them in their respective FXP IPs, shows that the use of 32 bit FXP IPs lead to the same performance. The use of 16 bit FXP IPs increases the error from 4% to 6%.

Figure S7b and c investigate how the error develops if 16 bit FXP IPs ($\gamma_1 = 1.0$ and $\gamma_2 = 1.0$) and 32 bit FXP IPs ($\gamma_1 = 1.0$ and $\gamma_2 = 1.0$) are used for the $H5$ and $HY$ layers (and the rest is still using 16 bit FXP IPs) too. This allows us to focus only on the effects on the error created by the discretization of the latent variables first. For learning, the latent variables are converted into 32 bit floating point numbers and then 32 bit FLP weights are updated based on this information. These 32 bit FLP weights are then converted in FXP weights for simulating the SbS network again. The result is that the error of the 32 bit FXPs is only slightly different to the 32 bit FLP IP variant. However, the 16 bit FXP IPs are having large errors for larger number of neurons in $H5$ (256, 512, and 1024). We are not sure why this is happening. Inspection of the learned weights matrices didn't reveal any obvious distortions. Figure S7a shows that already using 16 bit FXP numbers with known good weights and 1024 neurons leads to bad performance values. This might be a problem caused by the available dynamical range for small numbers. The SbS algorithm is multiplicative in nature. If a latent variable gets zero then it stays zero for the rest of the simulation. One possibility for the bad performance might be that the occuring fluctuations (by e.g. the stochastic spikes) drive latent variables into zeros just because of the quantization of numbers. Adding learning on top seems to strengthen the destructive effect even more.

Switching learning to their respective FXP update rules for the weights, as presented in figure S7c, reveals a roughly similar behavior for the 16 bit FXP case. In the case with 32 bit FXP something strange is happening. Learning seems to lead to overfitting after only a few learning steps (see figure S8). We couldn't find parameters that would prevent this overfitting. Figure S7c shows the performance of the 32 bit FXP after only 5 learning steps.

### 3 'RUN TIME' SIMULATION

For a comparison of the performance of a modern CPU with the FPGA simulation, we used simulations under C++11, heavily enhanced for Intel CPUs with the Intel Math Kernel Library (MKL 2019 Update 3) for all vector operations (i.e. cblas_sscal), cblas_sdot, cblas_saxpy, cblas_scopy, and vsMul in single (32bit) precision floating point numbers and measured the time using the high resolution clock from the std::chrono class. For keeping the measurement errors low, we always measured the time over one million spikes. For the FPGA, we show the performance for the 36bit floating point numbers. Since the problems with the Linux driver and data communication with the FPGA card made it impossible to reliably measure the time required for such a simulation, we calculated the required time from the measured number of clock cycles for an operation and the 125 MHz clock which are found to work with the FPGA firmware. Since the FPGA has a precise timing compared to a normal CPU these calculation lead to very good estimates for the required processing time.

The simulations were run on a system with an Intel Core i5-6600k with 3.8GHz in single core Turbo Boost mode and with 3.6GHz when all four cores are under load. Also we used a system with dual Intel Xeon E5-2640v3 (2x 8 cores) with 3.4GHz in single core and and 2.6GHz with multi cores. All RAM DIMM slots were filled with identical modules. Thus the double memory channel for the Intel i5 and the quad memory memory mode on the Intel Xeon were active. For using more than one thread on the CPUs we used std::thread from C++11. The source code of these simulations can be found in the supplemental materials.

#### 3.1 C++ source code

```
#include <iostream>
#include <fstream>
#include <stdint.h>
#include <ctime>
#include <ratio>
#include <chrono>
#include <random>
#include <math.h>
#include <string>
#include <thread>
#include <vector>

#include "mkl.h"

// ------------------------------------------------------------------------
// ------------------------------------------------------------------------
// ------------------------------------------------------------------------
// ------------------------------------------------------------------------

// Function for producing spikes from a h-vector H with N element and a random number
uint16_t Spike(float * H, uint16_t N, uint32_t RandomNumber){

  // A counter, used for counting.
  uint16_t Counter = 0;

  // A variable that will hold the acumulative sum over the h-vector
  float Sum = 0;
```

```
  // Converting the random number in a float with the range [0,1)
  float RandomNumber_s = float(RandomNumber)/float(0xFFFFFFFF);

  // Now we check all h-vector elements, except the last
  for (Counter = 0; Counter < (N-1); Counter ++){
    // The the actual h-vector value to the sum ...
    Sum += H[Counter];
    // ... and check if we found the spike, if the sum surpasses the random number
    if (RandomNumber_s <= Sum){
      // We found our spike
      return Counter;
    }
  }

  // If it wasn't one of the other h-vector elements, it must be the last one by definition
  return N-1;
};

// ------------------------------------------------------------------------
// ------------------------------------------------------------------------
// ------------------------------------------------------------------------
// ------------------------------------------------------------------------

void Measure_RandomNumberGenerator(uint32_t TheSeed, uint32_t Repeat_Counter_Max, uint32_t CoreMax,
    uint16_t Parameter_A){

  // Variables for measuring time in high-resolution
  std::chrono::high_resolution_clock::time_point t1;
  std::chrono::high_resolution_clock::time_point t2;
  std::chrono::duration<double> time_span;
  double Time = 0.0;

  // Mersene Twister Random Generator MT19937
  std::mt19937 mt_rand(TheSeed);
  uint32_t RandomNumber = 0;

  // Number of spikes generates
  uint32_t NumberOfSpikes = 1000000;
  // Counter for the spike generation
  uint32_t SpikeCounter = 0;
  // Result of the spike generation process
  uint16_t SpikeID = 0;

  // The length of the used part of the h-vector
  uint16_t N_S = Parameter_A;

  // Norm used for normalizing the selected part of the h-vector (Temporary variable)
  float Norm = 0;

  // Test h-vector with the full length
  std::vector<float> P(N_S);
  std::vector<float> Ones(N_S);

  // Well, some profane counter. Used for counting in a for loop
```

```
uint32_t i = 0;

// Generate the filename for the log file with the results
std::string Filename("Random_" + std::to_string(TheSeed) +"of" + std::to_string(CoreMax) + "Para" +
  std::to_string(N_S)  + ".txt");

// Log file (kill the content on opening)
std::ofstream LogFile;
LogFile.open(Filename);

// Counter for repeating the tests
uint32_t Repeat_Counter = 0;

// Filling the one vector with ones
for (i = 0; i < N_S; i ++){
  Ones[i] = 1.0;
}

// We will repeat the test several times for a better statistic
// with different h-vectors and hoping to average out fluctuations
// from the OS
for (Repeat_Counter = 0; Repeat_Counter < Repeat_Counter_Max; Repeat_Counter++) {

  // Filling the h-vector with random number from [0,1) range
  // I want to reuse the same vector for different sizes
  // There are many of these random vectors we will average over later
  for (i = 0; i < N_S; i ++){
    P[i] = float(mt_rand())/float(0xFFFFFFFF);
  }

  // Normalize P
  Norm = cblas_sdot(N_S, Ones.data(), 1, P.data(), 1);
  cblas_sscal(N_S, (float)(1.0/(float)Norm), P.data(), 1);

  // Here starts the measurement of time
  t1 = std::chrono::high_resolution_clock::now();

  // This loop goes over the number of spikes
  for (SpikeCounter = 0; SpikeCounter < NumberOfSpikes; SpikeCounter ++) {
    // Draw a random number
    RandomNumber = mt_rand();
    // Generate a spike from the random number and the h-vector
    SpikeID = Spike(P.data(), N_S, RandomNumber);
  }

  // Here ends the measurement of time
  t2 = std::chrono::high_resolution_clock::now();

  // Calculating how much time has passed
  time_span = std::chrono::duration_cast<std::chrono::duration<double>>(t2 - t1);
  // Converting it into a double
  Time = time_span.count();

  LogFile << Repeat_Counter << " " << " " << Time << "\n";
```

```
    }

    // Closing the file with the results
    LogFile.close();

    return;
};

// ---------------------------------------------------------------------------------
// ---------------------------------------------------------------------------------
// ---------------------------------------------------------------------------------
// ---------------------------------------------------------------------------------

void Measure_H_Update(uint32_t TheSeed, uint32_t Repeat_Counter_Max, uint32_t CoreMax, uint16_t
    Parameter_A , uint16_t Parameter_B){

    // Variables for measuring time in high-resolution
    std::chrono::high_resolution_clock::time_point t1;
    std::chrono::high_resolution_clock::time_point t2;
    std::chrono::duration<double> time_span;
    double Time = 0.0;

    // Selected length for the input vector and the h vector
    uint32_t N_S = Parameter_A;
    uint32_t N_H = Parameter_B;

    // Counter, for the for loops
    uint32_t Counter = 0;

    // Mersene Twister Random Generator MT19937
    std::mt19937 mt_rand(TheSeed);
    uint32_t RandomNumber = 0;

    // Arrays and matrices for the SbS simulation
    std::vector<float> Ones(N_S*N_H);

    std::vector<float> V_Temp(N_S*N_H);

    std::vector<float> P(N_S);
    std::vector<float> H(N_H);
    std::vector<float> W(N_S*N_H);

    float Norm = 0.0;
    float * Temp_Pointer = NULL;

    // Epsilon constant for the SbS dynamics
    float Epsilon = 0.1;
    float OneDivOnePlusEps = 1.0 / (1.0 + Epsilon);

    // Number of spikes generates
    uint32_t NumberOfSpikes = 1000000;
    // Counter for the spike generation
    uint32_t SpikeCounter = 0;
```

```
// Result of the spike generation process
uint16_t SpikeID = 0;

// Counter for repeating the tests
uint32_t Repeat_Counter = 0;

// Generate the filename for the log file with the results
std::string Filename("SbS_" + std::to_string(TheSeed) +"of" + std::to_string(CoreMax) + "Para" + std::
  to_string(N_S) + "_" + std::to_string(N_H)  + ".txt");

// Log file (kill the content on opening)
std::ofstream LogFile;
LogFile.open(Filename);

// Pre-filling Ones with 1.0
for (Counter = 0; Counter < (N_S*N_H); Counter ++){
  Ones[Counter] = 1.0;
}

// We will repeat the test several times for a better statistic
// with different h-vectors and hoping to average out fluctuations
// from the OS
for (Repeat_Counter = 0; Repeat_Counter < Repeat_Counter_Max; Repeat_Counter++) {

  // Pre-filling P_Full with random numbers
  for (Counter = 0; Counter < N_S; Counter ++){
    P[Counter] = float(mt_rand())/float(0xFFFFFFFF);
  }
  // Normalize P
  Norm = cblas_sdot(N_S, Ones.data(), 1, P.data(), 1);
  cblas_sscal(N_S, (float)(1.0/(float)Norm), P.data(), 1);

  // Pre-filling W_Full with random numbers
  for (Counter = 0; Counter < (N_S*N_H); Counter ++){
    W[Counter] = float(mt_rand())/float(0xFFFFFFFF);
  }
  // Normalization of W
  for (Counter = 0; Counter < N_H; Counter ++){
    Temp_Pointer = (W.data()+Counter*N_S);
    Norm = cblas_sdot(N_S, Ones.data(), 1, Temp_Pointer, 1);
    cblas_sscal(N_S, (float)(1.0/(float)Norm), Temp_Pointer, 1);
  }

  // Pre-filling W_Full with 1
  for (Counter = 0; Counter < N_H; Counter ++){
    H[Counter] = 1.0;
  }
  // Normalize H via 1.0/N_H
  cblas_sscal(N_H, (float)(1.0/(float)N_H), H.data(), 1);

  // Here starts the measurement of time
  t1 = std::chrono::high_resolution_clock::now();

  // This loop goes over the number of spikes
```

```
    for (SpikeCounter = 0; SpikeCounter < NumberOfSpikes; SpikeCounter ++) {
      // Draw a random number
      RandomNumber = mt_rand();

      // Generate a spike from the random number and the h-vector
      SpikeID = Spike(P.data(), N_S, RandomNumber);

      // Copy the part of W which was selected by the spike
      // V_Temp = W(Spike,:)';
      Temp_Pointer = (W.data()+SpikeID);
      cblas_scopy(N_H, Temp_Pointer, N_S, V_Temp.data(), 1);

      // Element wise vector multiplication between the selected weights and H
      // V_Temp = V_Temp .* H
      vsMul(N_H,H.data(), V_Temp.data(),V_Temp.data());

      // Calculate the sum over V_Temp
      // sum(V_Temp)
      Norm = cblas_sdot(N_H, Ones.data(), 1, V_Temp.data(), 1);

      // H = H + V_Temp * Epsilon/Norm
      cblas_saxpy(N_H, Epsilon/Norm, V_Temp.data(),1, H.data(), 1);

      // Normalize H
      // H = H * (1/(1+epsilon))
      cblas_sscal(N_H, OneDivOnePlusEps, H.data(), 1);
    }

    // Here ends the measurement of time
    t2 = std::chrono::high_resolution_clock::now();

    // Calculating how much time has passed
    time_span = std::chrono::duration_cast<std::chrono::duration<double>>(t2 - t1);
    // Converting it into a double
    Time = time_span.count();

    LogFile << Repeat_Counter << " " << " " << Time << "\n";

  }

  // Closing the file with the results
  LogFile.close();

  return;
};

// ------------------------------------------------------------------------------
// ------------------------------------------------------------------------------
// ------------------------------------------------------------------------------
// ------------------------------------------------------------------------------


void Measure_OnlineLearning(uint32_t TheSeed, uint32_t Repeat_Counter_Max, uint32_t CoreMax, uint16_t
    Parameter_A , uint16_t Parameter_B){
```

```
// Variables for measuring time in high-resolution
std::chrono::high_resolution_clock::time_point t1;
std::chrono::high_resolution_clock::time_point t2;
std::chrono::duration<double> time_span;
double Time = 0.0;

// Selected length for the input vector and the h vector
uint32_t N_S = Parameter_A;
uint32_t N_H = Parameter_B;

// Counter, for the for loops
uint32_t Counter = 0;

// Mersene Twister Random Generator MT19937
std::mt19937 mt_rand(TheSeed);
uint32_t RandomNumber = 0;

// Arrays and matrices for the SbS simulation
std::vector<float> Ones(N_S*N_H);

std::vector<float> V_Temp(N_S*N_H);

std::vector<float> P(N_S);
std::vector<float> H(N_H);
std::vector<float> W(N_S*N_H);

float Norm = 0.0;
float * Temp_Pointer = NULL;

// Epsilon constant for the SbS dynamics
float Epsilon = 0.1;
float OneDivOnePlusEps = 1.0 / (1.0 + Epsilon);

// Learning rate
float Gamma = 0.0001;

// Offset rate for the weights
float Offset = 0.05;
float FactorA = Offset / (float)N_S;
float FactorB = 1.0 / (1.0 + Offset);

// Number of spikes generates
uint32_t NumberOfSpikes = 1000000;
// Counter for the spike generation
uint32_t SpikeCounter = 0;
// Result of the spike generation process
uint16_t SpikeID = 0;

// Counter for repeating the tests
uint32_t Repeat_Counter = 0;

// Generate the filename for the log file with the results
```

```
std::string Filename("Online_" + std::to_string(TheSeed) +"of" + std::to_string(CoreMax) + "Para" + std::to_string(N_S) + "_" + std::to_string(N_H)  + ".txt");

// Log file (kill the content on opening)
std::ofstream LogFile;
LogFile.open(Filename);

// Pre-filling Ones with 1.0
for (Counter = 0; Counter < (N_S*N_H); Counter ++){
  Ones[Counter] = 1.0;
}

// We will repeat the test several times for a better statistic
// with different h-vectors and hoping to average out fluctuations
// from the OS
for (Repeat_Counter = 0; Repeat_Counter < Repeat_Counter_Max; Repeat_Counter++) {

  // Pre-filling P_Full with random numbers
  for (Counter = 0; Counter < N_S; Counter ++){
    P[Counter] = float(mt_rand())/float(0xFFFFFFFF);
  }
  // Normalize P
  Norm = cblas_sdot(N_S, Ones.data(), 1, P.data(), 1);
  cblas_sscal(N_S, (float)(1.0/(float)Norm), P.data(), 1);

  // Pre-filling W_Full with random numbers
  for (Counter = 0; Counter < (N_S*N_H); Counter ++){
    W[Counter] = float(mt_rand())/float(0xFFFFFFFF);
  }
  // Normalization of W
  for (Counter = 0; Counter < N_H; Counter ++){
    Temp_Pointer = (W.data()+Counter*N_S);
    Norm = cblas_sdot(N_S, Ones.data(), 1, Temp_Pointer, 1);
    cblas_sscal(N_S, (float)(1.0/(float)Norm), Temp_Pointer, 1);
  }

  // Pre-filling W_Full with 1
  for (Counter = 0; Counter < N_H; Counter ++){
    H[Counter] = 1.0;
  }
  // Normalize H via 1.0/N_H
  cblas_sscal(N_H, (float)(1.0/(float)N_H), H.data(), 1);

  // Here starts the measurement of time
  t1 = std::chrono::high_resolution_clock::now();

  // This loop goes over the number of spikes
  for (SpikeCounter = 0; SpikeCounter < NumberOfSpikes; SpikeCounter ++) {

    // Adding a normalized offset to the weights
    cblas_saxpy(N_S*N_H, FactorA, Ones.data(), 1, W.data(), 1);
    cblas_sscal(N_S*N_H, FactorB, W.data(), 1);

    // Draw a random number
```

```
      RandomNumber = mt_rand();

      // Generate a spike from the random number and the h-vector
      SpikeID = Spike(P.data(), N_S, RandomNumber);

      // Copy the part of W which was selected by the spike
      // V_Temp = W(Spike,:)';
      Temp_Pointer = (W.data()+SpikeID);
      cblas_scopy(N_H, Temp_Pointer, N_S, V_Temp.data(), 1);

      // Element wise vector multiplication between the selected weights and H
      // V_Temp = V_Temp .* H
      vsMul(N_H,H.data(), V_Temp.data(),V_Temp.data());

      // Calculate the sum over V_Temp
      // sum(V_Temp)
      Norm = cblas_sdot(N_H, Ones.data(), 1, V_Temp.data(), 1);

      // H = H + V_Temp * Epsilon/Norm
      cblas_saxpy(N_H, Epsilon/Norm, V_Temp.data(),1, H.data(), 1);

      // Normalize H
      // H = H * (1/(1+epsilon))
      cblas_sscal(N_H, OneDivOnePlusEps, H.data(), 1);

      // W = W + V_Temp * Gamma/Norm
      cblas_saxpy(N_H, Gamma/Norm, V_Temp.data(),1, Temp_Pointer , N_S);

      // Normalization of updated W
      for (Counter = 0; Counter < N_H; Counter ++){
        Temp_Pointer = (W.data()+Counter*N_S);
        Norm = 1.0 / (1.0 + (Gamma/Norm) * V_Temp[Counter]);
        cblas_sscal(N_S, Norm, Temp_Pointer, 1);
      }

    }

    // Here ends the measurement of time
    t2 = std::chrono::high_resolution_clock::now();

    // Calculating how much time has passed
    time_span = std::chrono::duration_cast<std::chrono::duration<double>>(t2 - t1);
    // Converting it into a double
    Time = time_span.count();

    LogFile << Repeat_Counter << " " << " " << Time << "\n";

  }

  // Closing the file with the results
  LogFile.close();

  return;
};
```

```
// ------------------------------------------------------------------------
// ------------------------------------------------------------------------
// ------------------------------------------------------------------------
// ------------------------------------------------------------------------


int main(int argc, char** argv){

  // Parameters to the user interface
  uint16_t TestModus = 0;
  uint32_t Max_Thread = 1;
  uint32_t Repeat_Counter_Max = 1;
  uint16_t Parameter_A = 1024;
  uint16_t Parameter_B = 1024;

  // For the for loop over the threads
  uint32_t Counter_Thread = 0;


  // Handling the user input
  if ((argc != 6) && (argc != 5)){
    std::cout << "There is something wrong with the number of parameters.\n";
    std::cout << "Expected is\n";
    std::cout << "For test 0 (producing 1 million spikes):\n";
    std::cout << "./test 0 [Number of CPU Threads] [Number of repeatitions] [N_S]\n";
    std::cout << "For test 1 (1 million SbS updates):\n";
    std::cout << "./test 1 [Number of CPU Threads] [Number of repeatitions] [N_S] [N_H]\n";
    std::cout << "For test 2 (1 million SbS updates with online learning):\n";
    std::cout << "./test 2 [Number of CPU Threads] [Number of repeatitions] [N_S] [N_H]\n";
    std::cout << "-==- END -==-\n";
    return -1;
  }

  // Interpreting the user input
  TestModus = std::stoi(argv[1]);

  if ((TestModus == 0) && (argc == 6)){
    std::cout << "Too many parameters for test 0\n";
    std::cout << "For test 0 (producing 1 million spikes):\n";
    std::cout << "./test 0 [Number of CPU Threads] [Number of repeatitions] [N_S]\n";
    std::cout << "-==- END -==-\n";
    return -1;
  }

  Max_Thread = std::stoi(argv[2]);
  Repeat_Counter_Max = std::stoi(argv[3]);
  Parameter_A = std::stoi(argv[4]);
  if ((TestModus == 1) || (TestModus == 2)){
    Parameter_B = std::stoi(argv[5]);
  }

  // Sanitize the input
  switch(TestModus){
```

```
    case 0:
      std::cout << "Test: Generate 1 million spikes from a probability distribution.\n";
      break;

    case 1:
      std::cout << "Test: Process 1 million spikes with the SbS update rule.\n";
      break;

    case 2:
      std::cout << "Test: Process 1 million spikes with the SbS update rule and online weight updates.\n";
      break;

     default:
      std::cout << "Invalid test mode selected.\n";
      std::cout << "-==- END -==-\n";
      return -1;
      break;
  }

  std::cout << "Selected parameters are:\n";

  std::cout << "N_S: " << Parameter_A << "\n";
  if (Parameter_A <= 0){
    std::cout << "Invalid number for N_S\n";
    std::cout << "-==- END -==-\n";
    return -1;
  }
  if ((TestModus == 1) || (TestModus == 2)) {
    std::cout << "N_H: " << Parameter_B << "\n";
    if (Parameter_B <= 0){
      std::cout << "Invalid number for N_H\n";
      std::cout << "-==- END -==-\n";
      return -1;
    }

  }
  std::cout << "Number of CPU threads: " << Max_Thread << "\n";
  if (Max_Thread <= 0){
    std::cout << "Invalid number for CPU Threads\n";
    std::cout << "-==- END -==-\n";
    return -1;
  }

  std::cout << "Number of repetitions: " << Repeat_Counter_Max << "\n";
  if (Repeat_Counter_Max <= 0){
    std::cout << "Invalid number for repetitions\n";
    std::cout << "-==- END -==-\n";
    return -1;
  }

  // Let's create a collection of several threads
  std::vector<std::thread*> ThreadCollection(Max_Thread);
  for (Counter_Thread = 0; Counter_Thread < Max_Thread; Counter_Thread++){
```

```cpp
    ThreadCollection[Counter_Thread] = NULL;
  }

  std::cout << "-==- Test started -==-\n";

  // Now we start the threads
  switch (TestModus) {
    case 0:
      for (Counter_Thread = 0; Counter_Thread < Max_Thread; Counter_Thread++){
        ThreadCollection[Counter_Thread] = new std::thread(Measure_RandomNumberGenerator, Counter_Thread, Repeat_Counter_Max, Max_Thread, Parameter_A);
      }
      break;

    case 1:
      for (Counter_Thread = 0; Counter_Thread < Max_Thread; Counter_Thread++){
        ThreadCollection[Counter_Thread] = new std::thread(Measure_H_Update, Counter_Thread, Repeat_Counter_Max, Max_Thread, Parameter_A, Parameter_B);
      }
      break;

    case 2:
      for (Counter_Thread = 0; Counter_Thread < Max_Thread; Counter_Thread++){
        ThreadCollection[Counter_Thread] = new std::thread(Measure_OnlineLearning, Counter_Thread, Repeat_Counter_Max, Max_Thread, Parameter_A, Parameter_B);
      }
      break;
  }

  // Now we wait until the threads are gone
  for (Counter_Thread = 0; Counter_Thread < Max_Thread; Counter_Thread++){
    ThreadCollection[Counter_Thread]->join();
  }

  // Clearning up the threads
  for (Counter_Thread = 0; Counter_Thread < Max_Thread; Counter_Thread++){
    if (ThreadCollection[Counter_Thread] != NULL){
      delete ThreadCollection[Counter_Thread];
    }
    ThreadCollection[Counter_Thread] = NULL;
  }

  std::cout << "-==- Test finished -==-\n";

  return 0;
};
```

main.cpp
